# Loss of NPC1 enhances phagocytic uptake and impairs lipid trafficking in microglia

**DOI:** 10.1101/789511

**Authors:** Alessio Colombo, Lina Dinkel, Stephan A. Müller, Laura Sebastian Monasor, Martina Schifferer, Ludovico Cantuti-Castelvetri, Jasmin König, Lea Vidatic, Tatiana Bremova-Ertl, Silva Hecimovic, Mikael Simons, Stefan F. Lichtenthaler, Michael Strupp, Susanne A. Schneider, Sabina Tahirovic

**Affiliations:** German Center for Neurodegenerative Diseases (DZNE), Munich, Germany; Neuroproteomics, School of Medicine, Klinikum rechts der Isar, Technical University of Munich, Munich, Germany; Munich Cluster for Systems Neurology (SyNergy), Munich, Germany; Institute of Neuronal Cell Biology (TUM-NZB), Technical University of Munich; Faculty of Chemistry, Technical University Munich, Garching, Germany; Division of Molecular Medicine, Ruder Boskovic Institute, Zagreb, Croatia; Department of Neurology and German Center for Vertigo and Balance Disorders, Ludwig-Maximilians University, Munich, Germany; Department of Neurology, University Hospital Bern, Bern, Switzerland

**Keywords:** lipid trafficking, microglia, NPC, phagocytic impairment, proteome

## Abstract

Niemann-Pick type C disease is a rare neurodegenerative disorder mainly caused by mutations in *Npc1*, resulting in abnormal late endosomal/lysosomal lipid storage. Although microgliosis is a prominent pathological feature, consequences of NPC1 loss on microglial function remain uncharacterized. Here, we provide an in-depth characterization of microglial proteomic signatures and phenotypes in a NPC1-deficient (*Npc1*^*-/-*^) murine model and patient blood-derived macrophages. We demonstrate enhanced phagocytic uptake and impaired lipid trafficking in *Npc1*^*-/-*^ microglia that precede neuronal death. Loss of NPC1 compromises microglial developmental functions as revealed by increased synaptic pruning and deficient myelin turnover. Undigested myelin accumulates within multi-vesicular bodies of *Npc1*^*-/-*^ microglia while lysosomal degradation remains preserved. To translate our findings to human disease, we generated novel *ex vivo* assays using patient macrophages that displayed similar proteomic disease signatures and lipid trafficking defects as murine *Npc1*^*-/-*^ microglia. Thus, peripheral macrophages provide a novel promising clinical tool for monitoring disease progression and therapeutic efficacy in NPC patients. Our study underscores an essential role for NPC1 in immune cells and implies microglial therapeutic potential.

## Introduction

Niemann-Pick type C (NPC) disease is a rare lipid storage disorder (LSD) with heterogeneous presentations, including neurological, systemic and psychiatric symptoms. NPC patients suffer from ataxia, supranuclear saccade and gaze palsy, epileptic seizures, spasticity and progressive neurodegeneration, leading to premature death (Bonnot, Gama et al., 2019a, Patterson, Clayton et al., 2017, Vanier, 2010). Psychiatric symptoms comprise bipolar disorder, schizophrenia-like psychosis or major depression (Bonnot et al., 2019b). NPC mainly manifests during childhood, although juvenile and adult cases have also been reported (Bonnot et al., 2019a, Bonnot, Klunemann et al., 2019b, Garver, Francis et al., 2007, Shulman, David et al., 1995, Vanier, 2010). Approximately 95% of the patients carry autosomal recessive mutations in the *Npc1* gene, with the remaining cases being caused by mutations in the *Npc2* (Vanier, 2010). The *Npc1* gene encodes for a transmembrane and the *Npc2* for a soluble protein that are jointly responsible for the egress and recycling of lipoprotein-derived cholesterol from late endosomes/lysosomes towards other cell compartments like the endoplasmic reticulum (ER) or plasma membrane (Kwon, Abi-Mosleh et al., 2009, Li, Saha et al., 2016, Trinh, Brown et al., 2018). Impairment of this lipid trafficking route causes an abnormal accumulation of unesterified cholesterol and other lipids (e.g., glycosphingolipids, sphingomyelin and sphingosine) in late endosomal/lysosomal compartments, resulting in lysosomal dysfunction (Lloyd-Evans & Platt, 2010).

A mouse model that carries a spontaneous loss-of-function mutation within the *Npc1* gene (deletion of 11 out of its 13 transmembrane domains) is available (Loftus, Morris et al., 1997, Pentchev, Gal et al., 1980). This rodent model (BALB/cNctr-Npc1^m1N^/J, further abbreviated as *Npc1*^*-/-*^) reliably features early-onset human pathology, including neurodegeneration of vulnerable NPC regions such as the cerebellum, affecting in particular Purkinje cells, and the thalamus (Higashi, Murayama et al., 1993, Tanaka, Nakamura et al., 1988). The cortex and hippocampus appear less affected in NPC (Ong, Kumar et al., 2001). *Npc1*^*-/-*^ mice show first behavioral defects, such as mild cerebellar ataxia and tremor, at 6 weeks of age that become more prominent by 8 weeks. Severe ataxia, difficulties in food and water uptake and weight loss appear by 10-12 weeks of age (humane endpoint) (Smith, Wallom et al., 2009).

The molecular mechanism responsible for neuronal death in NPC is still not fully understood. It has been proposed that accumulation of lipids, particularly sphingosine, can induce an imbalance in calcium homeostasis and affect lysosomal trafficking (Lloyd-Evans, Morgan et al., 2008). Additionally, lipid accumulation within lysosomes and mitochondrial membranes may induce oxidative stress (Wos, Szczepanowska et al., 2016, Yu, Gong et al., 2005). Different studies linked NPC1 dysfunction to alterations in mammalian target of rapamycin complex 1 (mTORC1) and microtubule-associated proteins 1A/1B light chain 3B (LC3) signaling, suggesting that autophagy might be compromised in NPC (Castellano & Thelen, 2017, Ko, Milenkovic et al., 2005, Liao, Yao et al., 2007, Wos, Komiazyk et al., 2019). Although peripheral organs such as liver and spleen are affected by the disease, restoring NPC1 function in the central nervous system (CNS) prevents neurodegeneration and premature lethality of the *Npc1*^*-/-*^ mouse (Loftus, Erickson et al., 2002). However, restoring *Npc1* expression in neurons only does not fully rescue the phenotype and still results in lethality, suggesting that NPC1 is functionally important in other brain cells as well (Chen, Li et al., 2007, Lopez, Klein et al., 2011, Marshall, Watkins-Chow et al., 2018, Zhang, Strnatka et al., 2008). Noteworthy, the *Npc1* gene is ubiquitously expressed throughout the brain (Prasad, Fischer et al., 2000), with particularly high expression in oligodendrocytes and microglia (Zhang, Chen et al., 2014). Accordingly, it was shown that NPC1 function is needed for correct maturation of oligodendrocyte progenitor cells (OPCs) and the maintenance of the existing myelin (Takikita, Fukuda et al., 2004, Yu & Lieberman, 2013).

Microglia, as the resident immune cells of the CNS, regulate brain homeostasis by orchestrating essential physiological processes like myelination and synaptogenesis (Reemst, Noctor et al., 2016), but also actively contribute to pathophysiology of neurodegenerative disorders (Butovsky & Weiner, 2018, Keren-Shaul, Spinrad et al., 2017, Krasemann, Madore et al., 2017, Mathys, Adaikkan et al., 2017, Srinivasan, Friedman et al., 2016, Yeh, Hansen et al., 2017). Gene expression studies on Alzheimer’s disease (AD), amyotrophic lateral sclerosis (ALS), fronto-temporal lobar degeneration (FTLD) or multiple sclerosis (MS) have underscored microglial diversity and delineated homeostatic and disease signatures, often assigned as “microglial neurodegenerative phenotype” (MGnD) or “disease associated microglia” (DAM) (Butovsky & Weiner, 2018, Gotzl, Colombo et al., 2018, Keren-Shaul et al., 2017, Krasemann et al., 2017, Mazaheri, Snaidero et al., 2017, Safaiyan, Kannaiyan et al., 2016). Loss of *Npc1* function is also associated with a massive microgliosis (Baudry, Yao et al., 2003, Cologna, Cluzeau et al., 2014, Platt, Speak et al., 2016), and alterations of transcriptomic signatures have been reported in symptomatic *Npc1*^*-/-*^ mice (Cougnoux, Drummond et al., 2018). However, to which extent microglial activation plays a causative role in NPC pathology and merits therapeutic investigation is still debated (Lopez, Klein et al., 2012, Vance & Karten, 2014). While cell culture experiments suggested that *Npc1*^*-/-*^ microglia do not directly trigger neurotoxicity (Peake, Campenot et al., 2011) and microglial ablation in an NPC murine model was not beneficial (Gabande-Rodriguez, Perez-Canamas et al., 2018), additional studies supported beneficial effects of microglial modulation (Cougnoux et al., 2018, Smith et al., 2009, Williams, Wallom et al., 2014). Moreover, significant changes in inflammatory markers were reported in both pre-symptomatic murine model and human NPC cases, suggesting that immune response could be a precocious phenomenon preceding neuronal loss (Baudry et al., 2003, Cluzeau, Watkins-Chow et al., 2012, Cologna et al., 2014, Pressey, Smith et al., 2012). Taking into consideration that NPC pathology often manifests during childhood and brain development, early microglial dysregulation could have profound pathological consequences. Thus, deciphering microglial contribution to NPC neuropathology is of fundamental importance to judge whether modulation of the inflammatory response bears therapeutic potential.

In the present study we demonstrated that loss of NPC1 can lock microglia in a disease state, largely compromising their physiological functions. Additionally, we characterized a lipid trafficking defect in *Npc1*^*-/-*^ microglia and discovered that accumulation of undigested lipid material occurs primarily within multi-vesicular bodies (MVBs), while lysosomal degradation remains preserved. Furthermore, we demonstrated that macrophages of NPC patients recapitulate major defects observed in the brain, and thus present a valuable model for biomarker studies and clinical monitoring.

## Results

### Loss of NPC1 induces cholesterol storage in microglia and triggers disease associated proteomic and functional signatures

We confirm previous reports (Higashi et al., 1993, Tanaka et al., 1988) and showed that symptomatic *Npc1*^*-/-*^ mice (8 weeks) display a severe degeneration in the cerebellum, particularly affecting the Purkinje cell layer which was visualized by Calbindin immunostaining (Fig EV1A). Increased immunoreactivity of a myeloid-specific lysosomal marker CD68 (Fig EV1A) revealed microglial cells with amoeboid morphology (Cougnoux et al., 2018), suggesting their active phagocytic status (Daria et al., 2017, Walker & Lue, 2015, Zotova, Bharambe et al., 2013). However, regions like cortex (Fig EV1B) and hippocampus (Fig EV1C) also showed pronounced CD68 immunoreactivity in symptomatic *Npc1*^*-/-*^ mice, but without detectable alterations of neuronal marker NeuN, thus excluding overt neuronal loss.

*Npc1* is highly expressed in microglia (Zhang et al., 2014), implicating that it may have a cell autonomous function in immune cells. To test this hypothesis, we isolated primary cerebral microglia from symptomatic *Npc1*^*-/-*^ mice and cultured them *in vitro*. Using the cholesterol binding dye filipin (Friend & Bearer, 1981), we found an increased intracellular cholesterol content in *Npc1*^*-/-*^ microglia compared to WT (Fig 1A). Co-staining with CD68 showed that most of the cholesterol load in *Npc1*^*-/-*^ microglia was localized within late endosomes/lysosomes. Thus, cultured *Npc1*^*-/-*^ microglia isolated from the neurodegenerative brain environment still display a cholesterol storage phenotype. This result suggests that, in addition to neurons, NPC1 exerts a cell autonomous function in microglia and microglial phenotype should not be considered only as a bystander of neurodegeneration (Elrick et al., 2010, Lopez et al., 2011).

**Figure 1.**
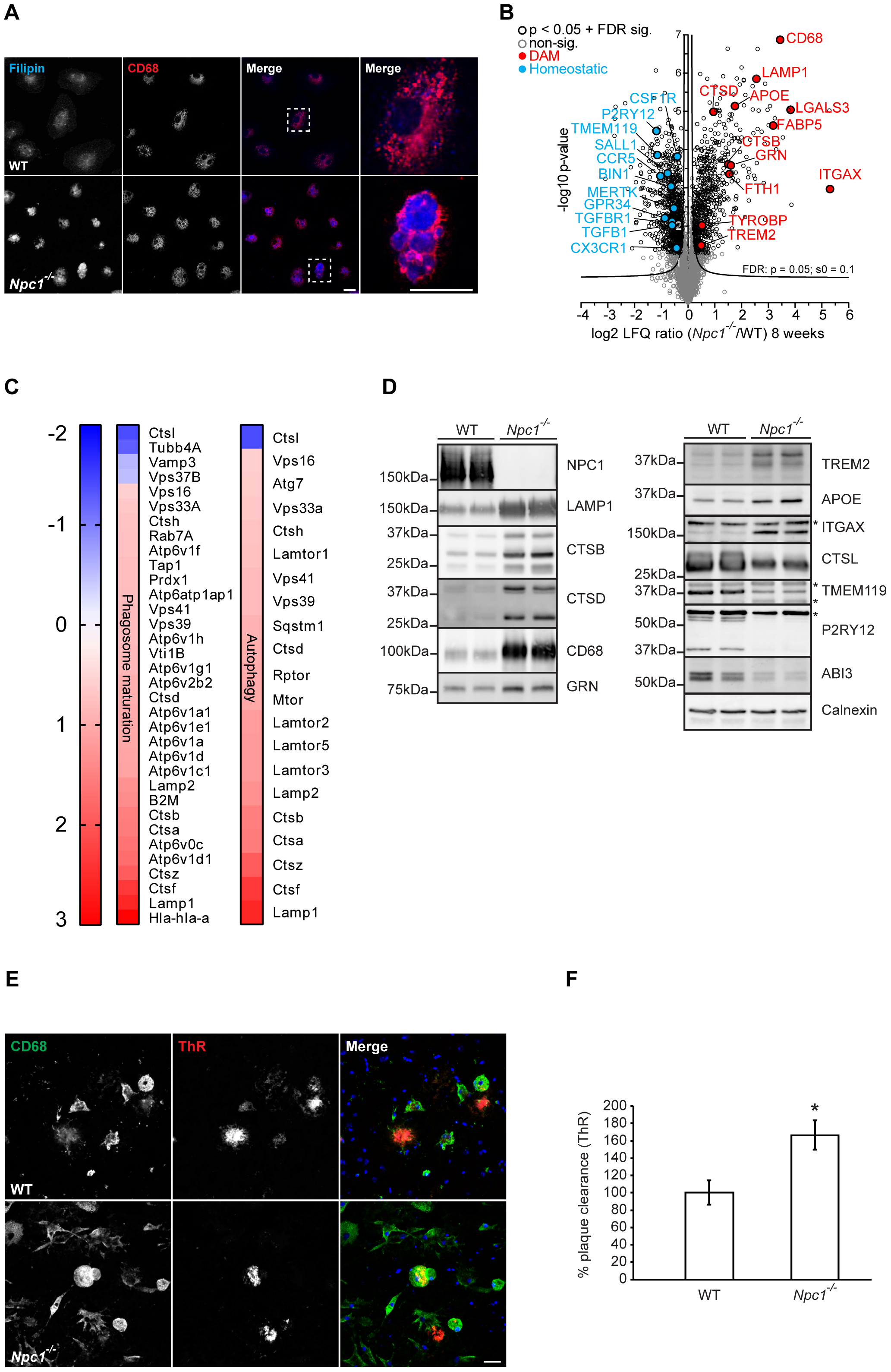
Loss of NPC1 induces microglial molecular and functional changes in symptomatic *Npc1*^*-/-*^ mice. **A** Immunocytochemistry of cultured primary microglia. Cholesterol content was visualized using Filipin dye (blue). *Npc1*^*-/-*^ cells show increased Filipin levels compared to WT, demonstrating cholesterol accumulation. Boxed regions are enlarged in right panels and show cholesterol accumulation within CD68 positive compartment (red) of *Npc1*^*-/-*^ microglia. Scale bars: 25 μm. **B** Proteome analysis of acutely isolated microglia from 8 weeks old *Npc1*^*-/-*^ and WT littermates reveals significantly decreased homeostatic and increased DAM markers in *Npc1*^*-/-*^ microglia. The negative log10 transformed p-value of each protein is plotted against its average log2 transformed LFQ ratio between *Npc1*^*-/-*^ and WT microglia. Significantly changed proteins are encircled in black. Downregulated homeostatic proteins are highlighted in blue and upregulated DAM proteins in red. Microglia was isolated and analyzed from 3 independent animals per group. **C** Phagosome maturation and autophagy are the most affected pathways upon NPC1 loss of function. The heatmap shows the average log2 transformed LFQ ratio between *Npc1*^*-/-*^ and WT microglia from significantly regulated proteins involved in phagosome maturation and autophagy according to IPA. Proteins related to mTOR signaling were manually added to the heatmap for autophagy. **D** Validation of MS data via western blot analysis. Representative immunoblots of acutely isolated *Npc1*^*-/-*^ and WT microglia show increased levels of DAM proteins (LAMP1, CTSB, CTSD, CD68, GRN, TREM2, APOE and ITGAX) and decreased levels of lysosomal protein CTSL and homeostatic signature markers (TMEM119, P2RY12 and ABI3). For each investigated protein, microglia isolated from 2 distinct mice per genotype were analyzed. Calnexin was used as loading control. * indicates unspecific bands. **E** Representative images of WT and *Npc1*^*-/-*^ microglia plated onto APPPS1 cryosection. Microglial lysosomes were visualized with an antibody against CD68 (green), while Aβ plaques were detected with Thiazine red (ThR) that stains fibrillar Aβ (red). Hoechst was used for nuclear staining (blue). Scale bar: 25 μm. **F** Quantification of Aβ plaque clearance was performed by comparing ThR positive area between a brain section incubated with microglia and the consecutive brain section where no cells were added. Values are expressed as percentages of amyloid plaque clearance normalized to the WT and represent mean ± SEM from 3 independent experiments (*p < 0.05, unpaired two-tailed Student’s t-test).

Recent findings in AD models revealed that under pathological conditions microglia can switch from a resting homeostatic to a DAM state that are characterized by distinct transcriptomic signatures (Krasemann et al., 2017). We profiled cerebral microglia acutely isolated from 8 weeks old *Npc1*^*-/-*^ and WT mice using an MS approach (Fig 1B, Table EV1). Quality control of microglia enriched fraction has been performed via western blot analysis using specific markers for microglia (Iba1), neurons (Tuj1), astrocytes (GFAP) and myelin (CNPase) (Fig EV2). Proteomic profiles of 8 weeks old microglia reveal major differences between *Npc1*^*-/-*^ and WT cells, featuring disease associated signatures (Fig 1B) (Keren-Shaul et al., 2017, Krasemann et al., 2017). Among significantly upregulated proteins, we identified TREM2, TYROBP, APOE, ITGAX and many late endosomal/lysosomal proteins including LAMP1/2, CD68, CTSB, CTSD and GRN. Among those, ITGAX showed the largest increase (55 fold). Concomitantly, markers associated with the homeostatic microglial function like P2RY12, TMEM119, CSF1R, CX3CR1, TGFB or TGFBR1 were decreased. Notably, IPA analysis of MS data revealed phagosome maturation and autophagy as the two pathways mostly affected by the loss of NPC1 (Fig 1C, Fig EV3), implicating a functional role of NPC1 in regulation of intracellular trafficking in microglia. Western blot analysis of selected homeostatic/DAM signature proteins (Fig 1D) and additional regulators of lysosome/autophagy pathways (Fig EV4) confirmed our MS data.

To study the functional consequences of altered microglial proteomic signatures, we used an *ex vivo* β-amyloid (Aβ) plaque clearance assay (Bard, Cannon et al., 2000, Claes, Van Den Daele et al., 2019, Xiang, Werner et al., 2016). To this end, microglial phagocytic clearance of Aβ plaques was determined by measuring fibrillar Aβ load visualized by Thiazine Red (ThR) staining (Fig 1E). Microglia from 8 weeks old *Npc1*^*-/-*^ mice showed a 1.6 fold higher phagocytic clearance of amyloid aggregates compared to WT cells (Fig 1F), suggesting that lysosomal degradation in *Npc1*^*-/-*^ microglia is functional.

Taken together, our data show that a prominent neuroinflammation in *Npc1*^*-/-*^ symptomatic mice is molecularly characterized by altered microglial proteomic signatures of intracellular trafficking, lysosomal function and phagocytosis and functionally reflected by increased phagocytic clearance.

### Early changes in microglial Npc1^-/-^ proteomic signatures occur prior to neuronal death and correlate with increased phagocytosis

Lack of NPC1 is likely to affect microglial function already during developmental stages when microglia are critically required for synaptic pruning and successful myelination that are pre-requisites for proper neuronal connectivity. To this end, we analyzed microglial phenotype at pre-symptomatic stages of *Npc1*^*-/-*^ mice (P7). In agreement with previous studies (Baudry et al., 2003, Ong et al., 2001, Tanaka et al., 1988), calbindin immunostaining delineated an intact layer of Purkinje cells, whereas cerebellar CD68 immunoreactivity was already upregulated in *Npc1*^*-/-*^ late endosomes/lysosomes when compared to WT cells (Fig 2A). This result implicates that despite preserved neuronal environment, cerebellar *Npc1*^*-/-*^ microglia were already activated at pre-symptomatic disease stages. Importantly, we also detected higher CD68 levels in less affected *Npc1*^*-/-*^ regions including cortex (Fig 2B) and the hippocampus (Fig EV5), similarly as shown for the symptomatic stages (Fig EV1).

**Figure 2.**
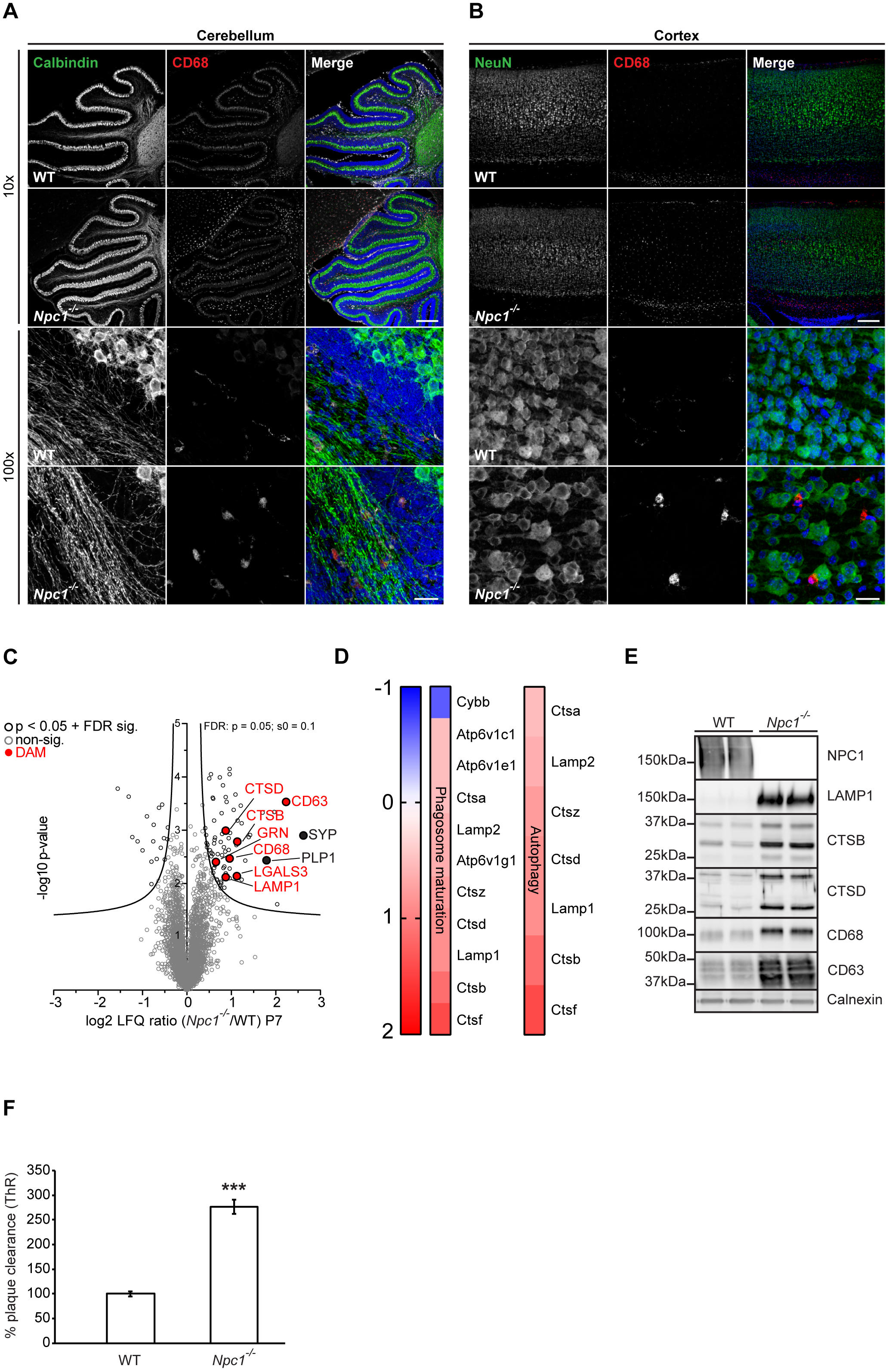
Switch in microglial proteomic signatures and increased phagocytosis precede neuronal loss in pre-symptomatic *Npc1*^*-/-*^ mice. **A-B** Immunostaining of cerebellum (**A**) and cortex (**B**) of WT and *Npc1*^*-/-*^ brain sections using antibodies against neuronal markers (green) calbindin (Purkinje cells, cerebellum), NeuN (cortex) and lysosomal microglial marker CD68 (red). At P7 no neurodegeneration was observed, neither in the cerebellum nor in the cortex of *Npc1*^*-/-*^ mice (10x, upper panels). In contrast, CD68 immunoreactivity showed activated *NPC1*^*-/-*^ microglia with amoeboid morphology already at this early pathological stage in both regions (100x, lower panels). Hoechst was used for nuclear staining (blue). Scale bars: 250 μm (10x, upper panels) and 25 μm (100x, lower panels). **C** Proteome analysis of acutely isolated microglia from P7 *Npc1*^*-/-*^ and WT littermates. The negative log10 transformed p-value of each protein is plotted against its average log2 transformed LFQ ratio between *Npc1*^*-/-*^ and WT microglia. Significantly changed proteins are encircled in black. Upregulated DAM proteins are highlighted in red and synaptic protein SYP and myelin PLP1 in black. Microglia were isolated and analyzed from 3 different animals per group. **D** *Npc1* deletion affects phagosome maturation and autophagy already at pre-symptomatic phase. The heatmap shows the average log2 transformed LFQ ratio between *Npc1*^*-/-*^ and WT microglia of all significantly regulated proteins involved in phagosome maturation and autophagy according to IPA. **E** Validation of MS microglia results via western blot analysis. Representative immunoblots of acutely isolated microglia lysates from *Npc1*^*-/-*^ mice and WT littermates showing increased level of lysosomal markers (LAMP1, CTSB, CTSD, CD68 and CD63). For each investigated protein, microglia isolated from 2 distinct pups per genotype have been analyzed. Calnexin was used as a loading control. **F** *Npc1*^*-/-*^ microglia from P7 pups already show increased phagocytic capacity towards Aβ. Quantification of Aβ plaque clearance was performed by comparing ThR positive area between a brain section incubated with microglia and the consecutive brain section where no cells were added. Values are expressed as percentages of amyloid plaque clearance normalized to the WT and represent mean ± SEM from 3 independent experiments (***p < 0.001, unpaired two-tailed Student’s t-test).

To pinpoint early microglial molecular alterations, we analyzed the proteome of cerebral microglia from pre-symptomatic *Npc1*^*-/-*^ animals. Although MS analysis of acutely isolated microglia from P7 *Npc1*^*-/-*^ mice showed less dramatic changes compared to the MS profile of 8 weeks old *Npc1*^*-/-*^ microglia (Fig 1B), most of the late endosomal/lysosomal markers, including LAMP1, CD63, CD68, CTSB, CTSD, GRN and others, were already significantly upregulated (Fig 2C, Table EV2). Upregulation of late endosomal/lysosomal proteins endorses microglial state switch as an early event in the NPC neuropathological cascade. Similar to symptomatic stage, canonical pathway analysis of microglia from pre-symptomatic stages highlighted autophagy and phagosome maturation as the two most significantly altered pathways (Fig 2D, Fig EV3). Several proteins found upregulated in the proteomic analysis (LAMP1, CTSB, CTSD, CD68 and CD63) were confirmed via western blot analysis (Fig 2E). The most significantly changed protein at the pre-symptomatic stage was the late endosomal and exosomal marker CD63 (Fig 2C and E, Table EV2), which suggests that defects in lipid trafficking and sorting may be among earliest pathological alterations in *Npc1*^*-/-*^ microglia (Kanerva, Uronen et al., 2013, Piper & Katzmann, 2007). Next, we tested whether proteomic fingerprint alterations were accompanied by functional changes in microglia. To this end, we performed the *ex vivo* Aβ plaque clearance assay. Microglia from pre-symptomatic *Npc1*^*-/-*^ mice showed more than 2.5-fold increased phagocytic clearance compared to WT microglia (Fig 2F). Thus, our data demonstrate that molecular and functional alterations of microglia are among earliest pathological hallmarks in NPC that occur independently from the neuronal loss observed at later stages.

### Npc1^-/-^ microglia display enhanced uptake but impaired turnover of myelin

It has been shown that *Npc1*^*-/-*^ mice exhibit reduced myelination (Weintraub, Abramovici et al., 1987, Weintraub, Abramovici et al., 1985) that has been linked to impairments in oligodendrocyte development (Qiao, Yang et al., 2018, Takikita et al., 2004, Yan, Lukas et al., 2011, Yu & Lieberman, 2013). However, it is well appreciated that microglia engulf and clear myelin debris, regulating thereby myelin turnover. This process can be impaired by cholesterol accumulation in microglia (Cantuti-Castelvetri, Fitzner et al., 2018, Schmitt, Castelvetri et al., 2015). To compare myelin levels in the *Npc1*^*-/-*^ and WT brain sections we used a co-staining for a myelin protein CNPase (2’,3’-cyclic-nucleotide 3’-phosphodiesterase) and compact myelin (Fluoromyelin) (Zhao, Tian et al., 2006) (Fig 3A). Symptomatic *Npc1*^*-/-*^ mice showed reduced levels of both CNPase and Fluoromyelin compared to WT (Fig 3A). Next, we analyzed removal of myelin debris by *Npc1*^*-/-*^ microglia. We found that in cerebellum (Fig 3B), cortex (Fig 3C) and the hippocampus (Fig EV6A) of symptomatic *Npc1*^*-/-*^ mice almost all CD68 positive cells accumulated Fluoromyelin intracellularly. Similarly, myelin accumulation was detected within microglia of symptomatic *Npc*-deficient mice (Gabande-Rodriguez, Perez-Canamas et al., 2018). Importantly, our proteomic analysis identified increased levels of a myelin specific protein (e.g., proteolipid protein 1, PLP1) and protein involved in microglial phagocytosis of myelin (e.g., galectin 3/LGALS3) in *Npc1*^*-/-*^ microglia already at pre-symptomatic stages (Fig 2C, Table EV2). Increased levels of PLP1 in *Npc1*^*-/-*^ microglia were also confirmed by western blot analysis (Fig EV6B). Taken together, our analysis suggests that *Npc1*^*-/-*^ hyper-reactive microglia accumulate myelin both at early pre-symptomatic as well as symptomatic stages and these changes are in line with observed proteomic alterations of microglia.

**Figure 3.**
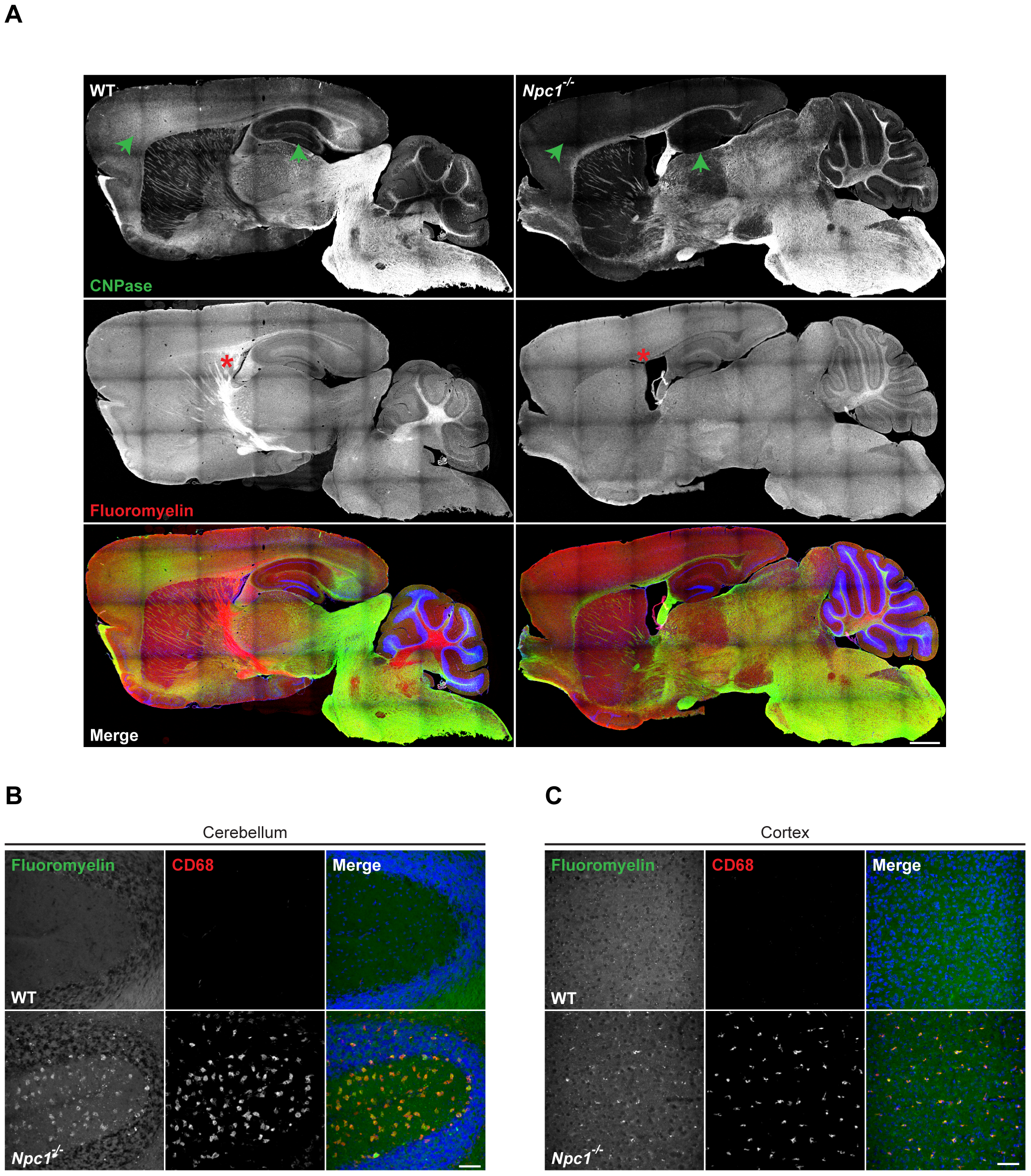
Hypomyelination phenotype and increased accumulation of myelin in microglia of symptomatic *Npc1*^*-/-*^ mice. **A** Tile scan of whole brain section from 8 weeks old WT and *Npc1*^*-/-*^ mice immunostained using antibody against CNPase as a general myelin marker (green) and Fluoromyelin dye that visualizes compact myelin (red). Hoechst was used for nuclear staining. Significantly reduced myelin levels are observed in *Npc1*^*-/-*^ cortex and hippocampus as shown by reduced CNPase immunoreactivity (green arrowheads). Fluoromyelin also reveals reduced levels of myelin in *Npc1*^*-/-*^ mouse compared to WT control (red star). Scale bar: 750 μm. **B-C** Microglia engulf myelin in symptomatic *Npc1*^*-/-*^ mice. Immunohistological analysis of 8 weeks old mouse brain sections demonstrates Fluoromyelin (green) positive myelin debris inside of CD68 positive (red) late endosomal/lysosomal compartments of *Npc1*^*-/-*^ cerebellum (**B**) and cortex (**C**) in contrast to WT. Scale bars: 50 μm

Increased levels of myelin proteins within *Npc1*^*-/-*^ microglia could reflect a higher uptake, but also possible defects in myelin turnover, or both. To address this question, we explored an *ex vivo* assay of myelin clearance (Fig EV7). Similarly, as we have described for the Aβ plaques, microglia from *Npc1*^*-/-*^ pre-symptomatic mice showed higher uptake of myelin, visualized by the increased levels of Fluoromyelin within CD68 positive cells compared to WT controls. However, instead of being degraded, the uptaken myelin accumulated within *Npc1*^*-/-*^ microglia (Fig EV7), suggesting possible impairments in myelin turnover.

Next, we investigated phagocytosis of exogenously added myelin in cultured *Npc1*^*-/-*^ microglia isolated from pre-symptomatic mice. As previously described (Cantuti-Castelvetri et al., 2018), cultured microglia have been pulsed for 6 h with purified and fluorescently labeled myelin and myelin turnover was monitored over 72 h (Fig 4A). After 6 h, myelin was mainly found inside of the CD68 positive late endosomal/lysosomal compartments in both WT and *Npc1*^*-/-*^ cells. At 48 h, most of the fluorescent labeling in WT cells was confined within vesicles outside of CD68 positive late endosomal/lysosomal compartment (Fig 4A and B). Immunostaining for the membrane marker Perilipin 2 confirmed that these vesicles were lipid droplets, cellular organelles specialized for lipid recycling and storage and crucial for cell metabolism (Olzmann & Carvalho, 2019) (Fig 4C). In contrast to WT cells, *Npc1*^*-/-*^ microglia accumulated labeled myelin within the late endosomal/lysosomal compartment, suggesting an impairment in myelin turnover and recycling into lipid droplets (Fig 4B). Even after 72 h, *Npc1*^*-/-*^ lysosomes still contained the engulfed myelin and completely lacked fluorescently labeled lipid droplets (Fig 4A), demonstrating a severe impairment and not just a delay in the myelin degradation process. To further confirm microglial impairment in myelin processing in *Npc1*^*-/-*^ microglia, cells at 72 h were analyzed for lipid droplet formation using the lipophilic dye Nile red (Greenspan, Mayer et al., 1985) (Fig EV8). Imaging analysis confirmed the presence of Nile red positive lipid droplets localized to the cell periphery in WT microglia. Again, in *Npc1*^*-/-*^ microglia lipid droplet staining could not be observed and Nile red visualized accumulated myelin within late endosomes/lysosomes.

**Figure 4.**
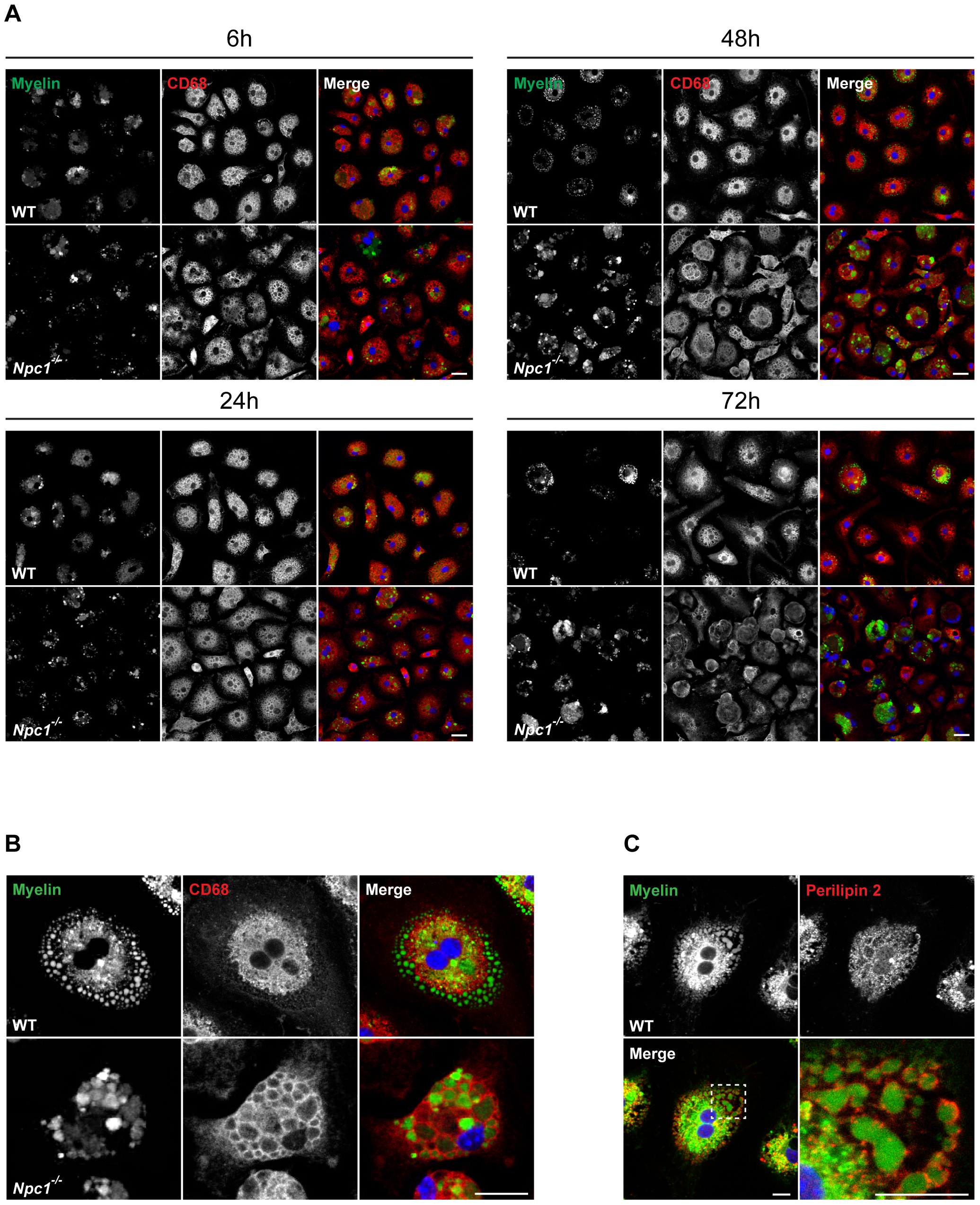
*Npc1*^*-/-*^ microglia display impairment in myelin turnover. **A-B** *In vitro* myelin phagocytosis assay. Cultured P7 primary microglia were incubated with fluorescently labeled myelin (green) and analyzed at 24, 48 and 72 h. Microglial lysosomes were stained with anti-CD68 antibody (red). Hoechst was used for nuclear staining (blue). **A** Both WT and *Npc1*^*-/-*^ microglia showed efficient myelin uptake at 6h. At 24 h (lower left panels) fluorescent labeling could be found within CD68 positive late endosomal/lysosomal compartments in both WT and *Npc1*^*-/-*^ microglia. At 48 h (upper right panels and higher magnification images shown in **B**), fluorescently labeled lipid vesicles were observed in WT microglia, indicating myelin turnover. In contrast, *Npc1*^*-/-*^ microglia accumulated myelin within the CD68 positive compartment and no fluorescently labeled lipid vesicles were detected. At 72 h (lower right panels), in most of the control cells myelin was degraded or signal could be detected in lipid vesicles. In *Npc1*^*-/-*^ microglia, myelin signal was still within the CD68 positive compartment, suggesting compromised myelin turnover. Scale bars: 25 μm. **C** Cultured WT primary microglia at 48 h were also stained against lipid droplet marker Perilipin 2 (red). Boxed region is enlarged in lower right panel showing that lipid vesicles forming after myelin turnover are lipid droplets. Hoechst was used for nuclear staining (blue). Scale bars: 10 μm.

### *Myelin accumulates in late endosomes/MVBs in NPC1*^*-/-*^ mice

To further characterize myelin accumulation, we performed an electron microscopy (EM) analysis of acutely isolated microglia from pre-symptomatic *Npc1*^*-/-*^ and WT mice. This analysis revealed pronounced accumulation of late endosomes/MVBs within *Npc1*^*-/-*^ microglia (Fig 5A and B (1) and (2)). Morphological analysis showed that most of these vesicles contained undigested myelin (Fig 5A and B (2)), further supporting that *Npc1*^*-/-*^ microglia hold an increased phagocytic capacity towards myelin, but fail in its processing. Interestingly, no obvious morphological alterations of lysosomes (Fig 5A) have been observed by EM. This is in agreement with myelin trafficking defect that occurs in late endosomes/MVBs, suggesting that myelin does not reach lysosomes where it is normally processed. To confirm myelin accumulation in late endosomes/MVBs, rather than lysosomes, we performed a pulse/chase experiment with primary microglia from pre-symptomatic *Npc1*^*-/-*^ mice (Fig 5C). Microglial cells were supplemented with green labeled myelin for 15 min, and myelin transport was followed over 1 h using fluorescently labeled CTX to visualize endosomes and DQ-BSA to label lysosomes. Confocal microscopy analysis revealed that in WT cells most of the phagocytosed myelin reached lysosomal compartment after 1 h. In contrast, myelin was poorly co-localizing with DQ-BSA in *Npc1*^*-/-*^ microglia (Fig 5D). Taken together, despite the enhanced phagocytic uptake by *Npc1*^*-/-*^ microglia, lipid trafficking towards the lysosomes was impaired, thus resulting in myelin accumulation in late endosomes/MVBs and defects in lipid clearance.

**Figure 5.**
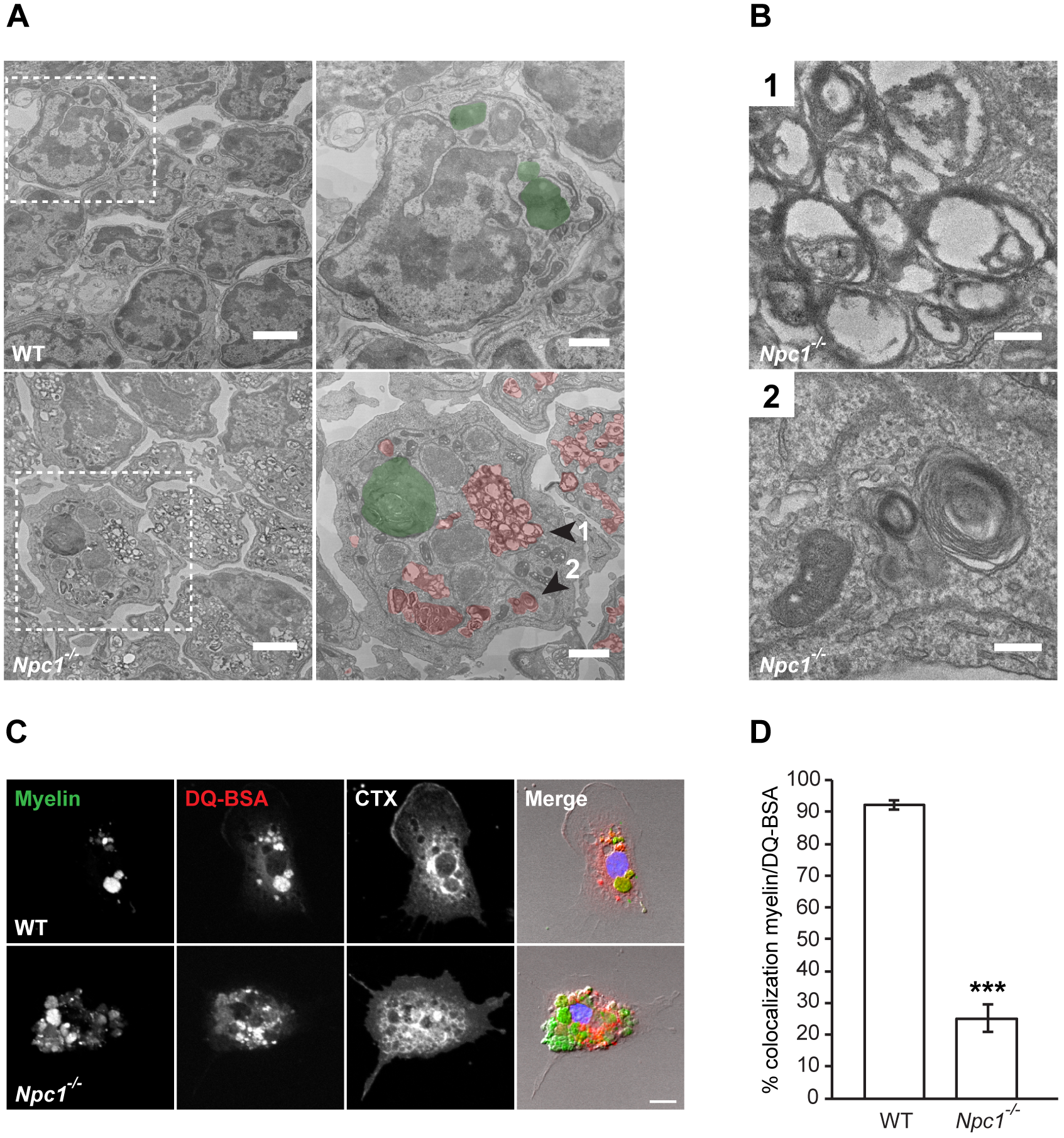
Myelin accumulation in late endosomes/MVBs of *Npc1*^*-/-*^ microglia. **A** EM analysis of microglia acutely isolated from WT and pre-symptomatic *Npc1*^*-/-*^ mice shows significant intracytoplasmic accumulation of MVBs in *Npc1*^*-/-*^ cells. Boxed regions are depicted at higher magnification in right panels (MVBs are pseudocolored in red). Scale bars: 2 μm (left panels) and 1 μm (right panels). **B** Representative high magnification images of *Npc1*^*-/-*^ cell shown in Fig. 1a (arrowheads 1 and 2) revealing accumulation of MVBs (1). Undigested myelin could be detected within MVBs (2). Scale bars: 0.2 μm. **C** Transport of phagocytosed myelin (green) was followed over 1h in P7 WT and *Npc1*^*-/-*^ microglia using DQ-BSA to visualize lysosomal compartment (red) and fluorescently labeled cholera toxin (CTX, white) as tracer to visualize endocytic vesicles. Scale bar: 10 μm. **D** Quantification of the *in vitro* myelin transport assay shows that, in contrast to WT control, phagocytosed myelin did not reach the lysosomal compartment in *Npc1*^*-/-*^ microglia. Values are expressed as percentages of myelin signal co-localizing with DQ-BSA positive lysosomes and represent mean ± SEM from 3 independent experiments (***p < 0.001, unpaired two-tailed Student’s t-test).

### Activated Npc1^-/-^ microglia display enhanced synaptic pruning during early brain development

Beyond its role in myelination, microglia has been shown to play a crucial role in shaping neuronal networks by synaptic pruning, particularly during early postnatal developmental (Reemst et al., 2016). Our results suggest that early postnatal defects detected in P7 *Npc1*^*-/-*^ microglia, reflected by disease associated proteomic signatures, could affect synaptic pruning. Indeed, by immunohistochemistry of brain sections from pre-symptomatic *Npc1*^*-/-*^ mice, we could readily detect the presence of the synaptic protein synaptophysin (SYP) inside of the CD68 positive late endosomes/lysosomes of *Npc1*^*-/-*^ microglia (Fig 6A) that could not be detected in WT microglia (Fig 6B). This result was endorsed by our proteome analysis of acutely isolated P7 microglia that revealed significantly increased levels of synaptophysin and other synaptic proteins in *Npc1*^*-/-*^ microglia (Fig 2C, Table EV2). Of note, our immunofluorescence analysis also revealed Hoechst positive signal inside of CD68 positive compartments of *Npc1*^*-/-*^ microglia which is indicative of cellular uptake by microglia (Fig 6A). Our data therefore provide evidence that altered microglial function caused by the loss of NPC1 may compromise neuronal connectivity and homeostasis via enhanced synaptic pruning.

**Figure 6.**
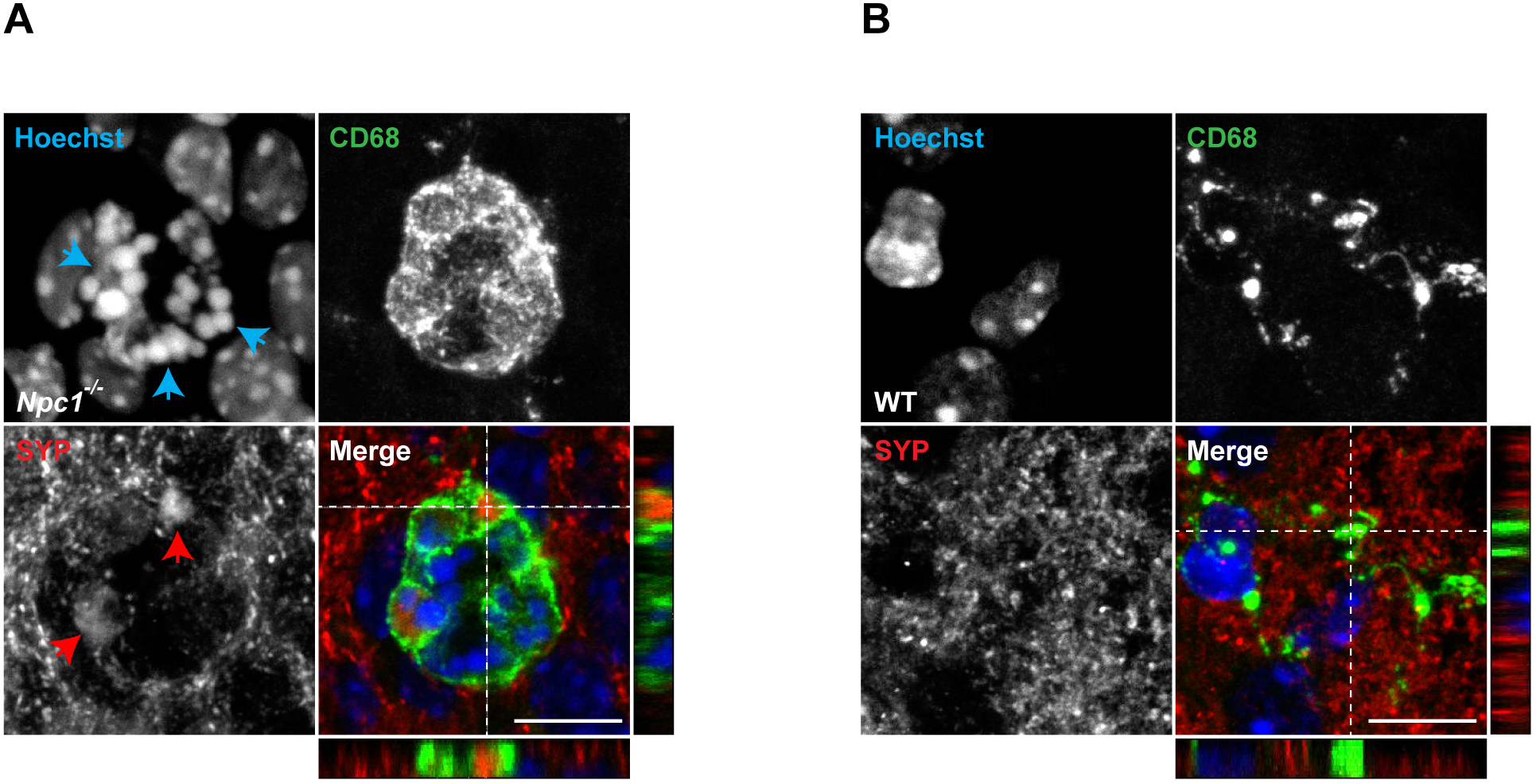
Chronically activated microglia engulf synapses in pre-symptomatic *Npc1*^*-/-*^ mice. **A-B** Immunostaining of *Npc1*^*-/-*^ (**A**) and WT (**B**) brain sections using antibodies against CD68 (green) and synaptic protein SYP (red). Hoechst was used for nuclear staining (blue). Representative image of a single *Npc1*^*-/-*^ microglial cell (**A**) shows condensed nuclear signal (blue arrowheads) and SYP positive material (red arrowheads) within a CD68 positive amoeboid microglia. In contrast, WT microglia (**B**) exhibit physiological CD68 punctate staining pattern (green) with no detectable nuclear or synaptic material within late endosomal/lysosomal compartment. Scale bars: 10 μm.

### Macrophages from NPC patients feature murine NPC1^-/-^ microglial phenotypes

*Npc1*^*-/-*^ microglia offer potential for therapeutic intervention and biomarker studies, but translation of animal models to human patients is limited due to the very small number of NPC patients (estimated at 1:100 000 live births) and limited access to viable human brain tissue (Hammond, Munkacsi et al., 2019). As a compromise, we explored human peripheral macrophages for their phenotypic alterations in NPC. First, we performed a proteomic analysis of peripheral blood monocyte-derived macrophages from two NPC patients and revealed increased levels of late endosomal/lysosomal proteins as observed in murine microglia (Table EV3). Indeed, proteins like LAMP1/2, CD63, CD68, ITGAX, CTSD, CTSB or LGALS3 showed higher levels in NPC patient macrophages compared to a healthy control (Fig 7A). As for the murine proteome, we further validated selected proteins by western blot analysis (Fig 7B). Strikingly, CD63 was still among the most significantly changed proteins upon loss of NPC1 (Table EV3) as we reported for murine microglia isolated from pre-symptomatic mice. This result is strengthening further the relevance of the lipid trafficking defect and MVB pathology in NPC.

**Figure 7.**
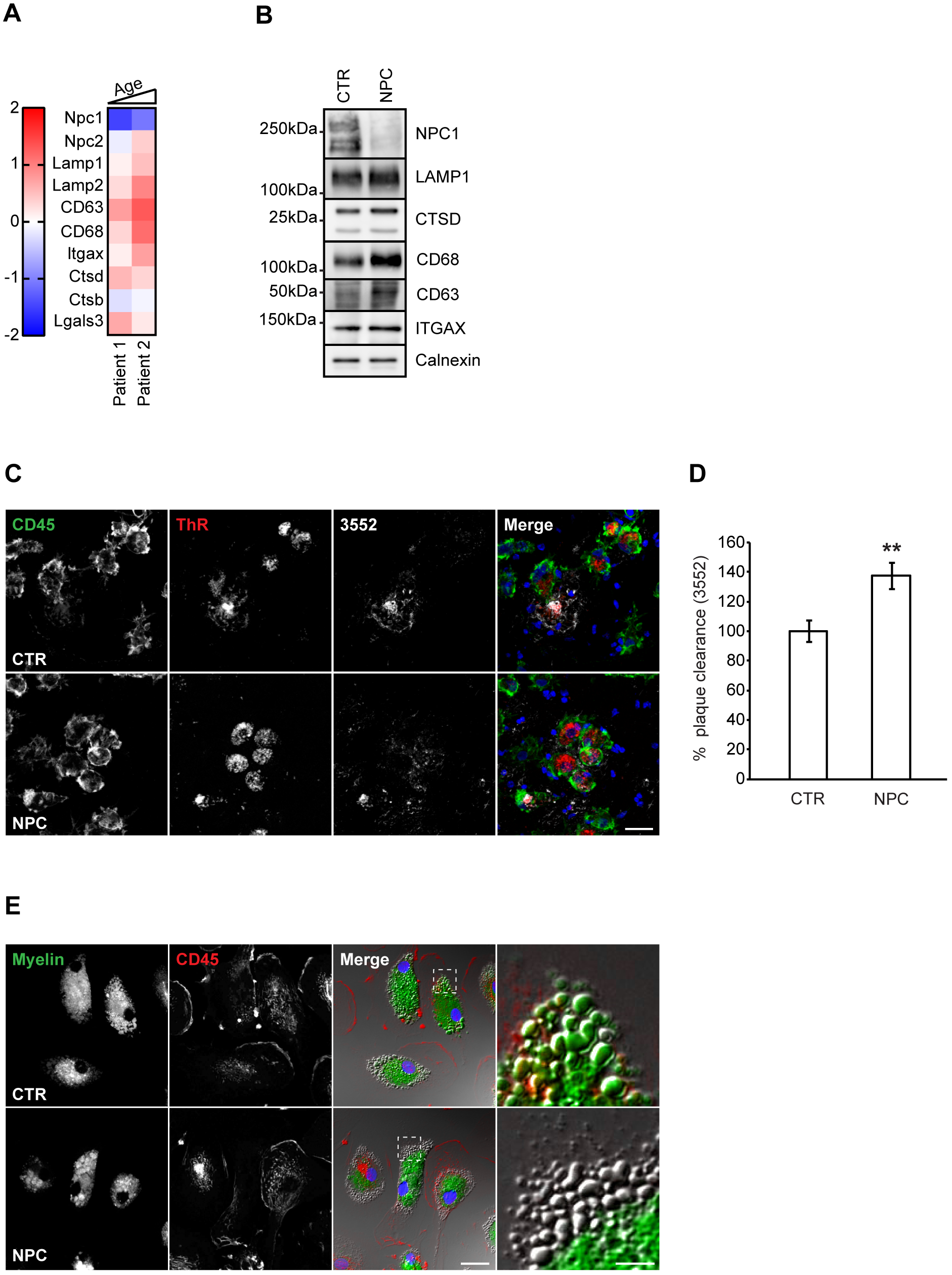
Human blood-derived macrophages from NPC patients resemble pathological alterations of murine *Npc1*^*-/-*^ microglia. **A** Heat map of selected proteins altered in NPC patients versus healthy controls based on proteomic analysis of macrophages. Noteworthy, an age dependent change was observed for some of the identified proteins (LAMP1/2, CD63, CD68, ITGAX or CTSB). **B** Validation of MS data via western blot analysis. Representative immunoblots reveal increased levels of late endosomal/lysosomal proteins in NPC patient-derived macrophages. Calnexin was used as loading control. **C** *Ex vivo* Aβ plaque clearance assay. Representative immunostaining showing macrophages (CD45, green) plated onto APPPS1 brain section that are clustering around and phagocytosing Aβ plaques, visualized with Thiazine red (ThR, plaque core, red) and 3552 anti-Aβ antibody (Aβ plaque, white). Scale bar: 25 μm. **D** Quantification of phagocytosed Aβ plaques. Values are expressed as percentages of amyloid plaque clearance normalized to the WT and represent mean ± SEM from 3 independent technical replicates (**p < 0.01, unpaired two-tailed Student’s t-test). **E** *In vitro* myelin phagocytosis assay. Human macrophages were fed with fluorescently labeled myelin (green) and analyzed after 48 h. Myeloid cells were visualized using an antibody against CD45 (red). Hoechst was used as nuclear staining (blue). Boxed regions are enlarged in right panels and show that human macrophages from healthy control efficiently uptake and turnover myelin as demonstrated by fluorescently labeled lipid droplets at the cell periphery. In contrast, we could not detect fluorescently labeled lipid droplets at the cell periphery in NPC patient-derived macrophages, suggesting trafficking defect that may preclude myelin turnover. Scale bars: 25 μm and 5 μm (enlargements).

Next, we functionally characterized patient macrophages using our *ex vivo* Aβ plaque clearance assay. NPC1-deficient human macrophages showed a higher amyloid plaque clearance capacity in comparison to control cells, resembling phenotypic features of rodent *Npc1*^*-/-*^ microglia (Fig 7C and D). Finally, to fully translate our observations from *Npc1*^*-/-*^ murine cells, we also performed the *in vitro* myelin phagocytic assay using patient cells. As we observed for the murine model, human NPC patient cells were capable to efficiently uptake exogenously added myelin, as judged by the intracellular fluorescent signal that could be detected in control and patient cells. At 48h, fluorescently labeled lipid droplets could be observed at the periphery in control cells. We again failed to detect fluorescently labeled lipid droplets at the cell periphery of NPC patient cells (Fig 7E), suggesting that trafficking defect may preclude myelin degradation and recycling into lipid droplets, as shown in murine *Npc1*^*-/-*^ microglia.

In conclusion, our data show that NPC patient macrophages recapitulate many of the key molecular and phenotypic features of murine *Npc1*^*-/-*^ microglia and thus represent valuable tools to identify biomarkers and monitor disease progression and therapeutic interventions in NPC patients.

## Discussion

Our study reveals pathological proteomic fingerprints of microglia in pre-symptomatic phase of NPC that associate with enhanced phagocytic uptake, increased synaptic pruning, impaired myelin turnover and its accumulation within MVBs. Microglia are responsible for the neuroinflammatory phenotype that occurs in brains of *Npc1*^*-/-*^ mice as infiltration of peripheral macrophages could not be detected in this model (Cho, Vardi et al., 2018, Cougnoux et al., 2018). However, the contribution of microglial activation to neurodegeneration in NPC is still being discussed (Baudry et al., 2003, Gabande-Rodriguez et al., 2018, Lopez et al., 2011, Peake et al., 2011, Pressey et al., 2012). In accordance with previous reports (Baudry et al., 2003, Cougnoux et al., 2018), we detected pronounced microglial reactivity throughout cortex and hippocampus prior to neuronal loss. Furthermore, our molecular and functional characterization supports microglial dysfunction as an early pathological insult in NPC.

In contrast to well characterized microglial transcriptomic profiles, proteomic signatures are largely unknown (Flowers, Bell-Temin et al., 2017, Rangaraju, Dammer et al., 2018, Sharma, Schmitt et al., 2015, Thygesen, Ilkjaer et al., 2018). Proteomic profiles of acutely isolated *Npc1*^*-/-*^ microglia revealed significant upregulation of numerous late endosomal/lysosomal proteins (e.g., LAMP1/2, CD63, CD68, CTSB and CTSD) already in pre-symptomatic mice. Some of the detected proteomic alterations were also identified in a recent transcriptomic study of microglia in symptomatic *Npc1*^*-/-*^ mice (Cougnoux et al., 2018). Importantly, our analysis of human macrophages from NPC patients revealed a myeloid dysfunction, resembling murine *Npc1*^*-/-*^ microglia (e.g., LAMP1/2, CD63, CD68, ITGAX or CTSD). Noteworthy, increased levels of Cathepsins were reported in brain and plasma of NPC patients (Alam, Getz et al., 2014, Cluzeau, Watkins-Chow et al., 2012). Along these lines, upregulation of pro-inflammatory gene expression in both murine and human brains as well as inflammatory CSF signatures of NPC patients suggest that immune changes are early pathological hallmarks of NPC (Cologna et al., 2014). Moreover, beside NPC, neuroinflammation is emerging as a common pathological denominator in other LSDs, suggesting that microglia are vulnerable to lysosomal impairments and that neuroinflammation can contribute to neuronal death (Bosch & Kielian, 2015, Platt, Boland et al., 2012). These findings are in line with our study showing that NPC1 loss results in a strong pro-inflammatory phenotype and dysregulated microglia at early pre-symptomatic disease stage. Accordingly, pharmacological or genetic immune modulations have been shown to be beneficial and improved life expectancy in NPC murine models (Cougnoux et al., 2018, Smith et al., 2009, Williams et al., 2014).

It is currently unclear how DAM signatures across various neurodegenerative disease reflect microglial function and whether they are predictive of beneficial or detrimental responses. We show here that DAM signatures correlate with increased phagocytic uptake both in murine *Npc1*^*-/-*^ microglia and in human NPC macrophages. Similarly, in LSD caused by granulin loss of function (*Grn*^*-/-*^) (Gotzl et al., 2018, Gotzl, Mori et al., 2014), DAM signatures correlated with increased microglial phagocytosis (Gotzl, Brendel et al., 2019). Surprisingly, microglial cells in AD acquire DAM signatures (Butovsky & Weiner, 2018, Krasemann et al., 2017), but display impaired phagocytic function (Daria et al., 2017, Hickman, Allison et al., 2008, Krabbe, Halle et al., 2013). Since AD and NPC microglia share many DAM markers, but still differ in phagocytic capacity towards Aβ, our data strengthen the fundamental importance of combining molecular fingerprint analysis with functional studies to identify molecular targets that are predictive of microglial function and could be explored for repair. NPC patients typically do not show amyloid plaque deposition, but do have increased Aβ production (Causevic, Dominko et al., 2018, Malnar, Hecimovic et al., 2014, Malnar, Kosicek et al., 2010, Yamazaki, Chang et al., 2001). This was ascribed as a consequence of the limited life expectancy of NPC patients, but it is also tempting to speculate that increased phagocytic Aβ clearance in NPC may compensate for the increased Aβ generation and thereby counteract amyloid deposition.

Uncontrolled and overt microglial activation during early *Npc1*^*-/-*^ development can compromise physiological processes and neuronal homeostasis. It is known that microglia play a role in synaptic pruning (Schafer, Lehrman et al., 2012, Stevens, Allen et al., 2007) and synaptic changes together with axonal pathology are reported in *Npc1*^*-/-*^ mice (Pressey et al., 2012). Of note, our data reveal an increased synaptic pruning in *Npc1*^*-/-*^ brains that could compromise neuronal connectivity as described in AD, schizophrenia and *Grn*^*-/-*^ models (Filipello, Morini et al., 2018, Hong, Beja-Glasser et al., 2016, Lui, Zhang et al., 2016, Sellgren, Gracias et al., 2019). Psychiatric symptoms have been described in NPC patients, including mental retardation, behavioral problems, schizophrenia-like psychosis, bipolar disorder, and attention deficit hyperactivity disorder (Bonnot et al., 2019b, Evans & Hendriksz, 2017). Further work is needed to clarify whether excessive microglial activity and synaptic pruning contribute to psychiatric manifestations reported in NPC patients. In addition to synaptic pruning, microglia orchestrate myelin homeostasis, especially during development. Microglia control OPC recruitment and differentiation (Domingues, Portugal et al., 2016, Miron, 2017) and are responsible for the clearance of myelin debris. Impaired removal of myelin debris by microglia can heavily compromise re-myelination upon injury (Cantuti-Castelvetri et al., 2018, Gabande-Rodriguez et al., 2018). Hypomyelination phenotype described in NPC mouse models and human patients has been mainly associated to a cell autonomous impairment of oligodendrocytes (Kodachi, Matsumoto et al., 2017, Qiao et al., 2018, Takikita et al., 2004, Weintraub et al., 1987, Weintraub et al., 1985, Yu & Lieberman, 2013). Our data reveal that at pre-symptomatic stage, when myelination process just started (Foran & Peterson, 1992), *Npc1*^*-/-*^ microglia already show increased myelin uptake. Furthermore, LGALS3, that is expressed at high levels in microglia, triggers OPC differentiation and, together with TREM2, regulates myelin phagocytosis (Thomas & Pasquini, 2018). Our proteomic dataset showed that LGALS3 level is strongly increased in *Npc1*^*-/-*^ microglia, further supporting an increased phagocytic capacity towards myelin. Accordingly, early accumulation of LGALS3 was found in serum of NPC patients, suggesting its potential as a biomarker for therapeutic monitoring (Cluzeau et al., 2012). This collective evidence suggests that, in addition to oligodendrocytes, hyperactive microglia may also contribute to aberrant myelination in NPC.

Despite efficient uptake, *Npc1*^*-/-*^ cells failed to process phagocytosed myelin. The observed defects in degradation of myelin in *Npc1*^*-/-*^ microglia and NPC patient macrophages are indeed in contrast to what has been observed for Aβ substrate where phagocytosed material was cleared more efficiently. Along these lines, it has been proposed that lysosomal function is preserved in NPC patient cells (Sarkar, Carroll et al., 2013, Schwerd, Pandey et al., 2017). Thus, we hypothesize that myelin degradation is impaired because phagocytosed material accumulates within late endosomes/MVBs that do not fuse with lysosomes. Impaired autophagy, vesicular fusion and MVB accumulation has been demonstrated in NPC (Ko et al., 2005, Liao et al., 2007, Phillips et al., 2008, Tharkeshwar, Trekker et al., 2017). Aβ substrate may be following an alternative route to reach the lysosome and thus being efficiently degraded. This observation is of importance because it suggests that the main lipid storage burden in NPC may not be occurring in lysosomes, but rather in late endosomal compartments/MVBs and therefore the term lysosomal storage disease may be revisited. Myelin accumulation that we observed in *Npc1*^*-/-*^ microglia may be occurring in CD63 positive intracellular compartment as this MVB/exosome marker is the most significantly altered protein in NPC patient macrophages and *Npc1*^*-/-*^ microglia. Thus, CD63 may be an early pathological marker in myeloid cells that could be explored for monitoring of therapeutic efficacy in NPC patients. Along these lines, another MVB protein (CHM2A) is also upregulated in pre-symptomatic phase of NPC, strengthening the relevance of MVB function for early pathogenesis of NPC (Cologna, Jiang et al., 2012)

Lipids deriving from myelin degradation are normally stored in droplets (Olzmann & Carvalho, 2019) that play a key regulatory function in cellular metabolism (Olzmann & Carvalho, 2019). Beyond their important metabolic role, lipid droplets can also prevent lipo-toxicity (Greenberg, Coleman et al., 2011, Krahmer, Farese et al., 2013) or protect cells from high reactive oxygen species (ROS) (Bailey, Koster et al., 2015, Liu, MacKenzie et al., 2017, Liu, Zhang et al., 2015), ER stress or mitochondrial damage (Nguyen, Louie et al., 2017, Olzmann & Carvalho, 2019, Torres, Balboa et al., 2017). Thus, disturbances in lipid droplet formation in *Npc1*^*-/-*^ microglia may contribute to increased cellular stress and neurotoxicity (Dionisio-Santos, Olschowka et al., 2019). Accordingly, increased ROS, cellular stress and mitochondrial dysfunction have all been associated with NPC (Yambire, Fernandez-Mosquera et al., 2019). Moreover, aberrant lipid droplet formation (Onal, Kutlu et al., 2017) may also be linked to autophagy defects described in NPC (Castellano & Thelen, 2017, Ko et al., 2005, Liao et al., 2007, Sarkar et al., 2013, Wos et al., 2019). Our proteomic analysis identified autophagosome formation and maturation as the most affected pathway in microglia upon loss of NPC1. Significant proteomic changes in proteins involved in autophagy further strengthen the regulatory role of NPC1 in intracellular trafficking (Ko et al., 2005, Liao et al., 2007, Sarkar et al., 2013, Schwerd et al., 2017). Alterations in mTOR regulators were also found in our macrophage proteome analysis, confirming that this defect occurs in NPC patient cells.

The increased lipid burden in *Npc1*^*-/-*^ microglia may contribute to further upregulation of late endosomal/lysosomal proteins in order to compensate for the lipid degradation failure. This negative feedback loop likely results in enhanced phagocytic uptake that further amplifies lipid burden and generates a chronic and self-sustained microglial activation that may be harmful to neurons. Thus, we believe that a pharmacological approach based on a combination of immune modulation and lipid reducing agents should be considered as a therapeutic strategy in NPC.

## Materials and Methods

### Animals

Male and female C57BL/6J (Jackson Laboratory stock N° 000664), BALB/cNctr-Npc1^m1N^/J (Jackson Laboratory stock N° 003092) (Loftus et al., 1997, Pentchev et al., 1980) and APPPS1 mice (Radde, Bolmont et al., 2006) were used in this study. Animals were group housed under specific pathogen-free conditions. Mice had access to water and standard mouse chow (Ssniff Ms-H, Ssniff Spezialdiäten GmbH, Soest, Germany) *ad libitum* and were kept in a 12/12-h light–dark cycle in IVC System. All experimental procedures were performed in accordance with the German animal welfare law and have been approved by the government of Upper Bavaria (license number ROB-55.2-2532.Vet_02-17-075).

### Human material

All studies were performed in accordance to the 1964 Declaration of Helsinki and were approved by the local Ethics Committee of the University of Munich. Two clinically affected *Npc1* mutation carriers (e.g., siblings, 17 and 24 years old males) and two healthy donors (24 years old female and 42 years old male) were included into this study. For LC-MS/MS analysis one healthy control has been used (42 years old male). All participants gave written informed consent prior to their inclusion in the study. NPC patients were from a Jordanian family with double consanguinity (e.g., the parents as well as the grandparents were first cousins). Disease onset was at around 10 years of age. Symptoms and disease course were typical of NPC with progressive ataxia, dystonia, dysarthria, abnormal eye movements with supranuclear gaze palsy, epilepsy, cognitive decline, sleep disturbances and autonomic involvement with bladder and gastrointestinal dysfunction. The older sibling is more severely affected; largely unresponsive, anarthric and wheelchair-dependent. The younger sibling can speak a few words and walk, but is very unstable, with frequent falls. The older sibling is being treated with anticholinergics and a benzodiazepine and the younger takes miglustat, baclofen, anticholinergics and a benzodiazepine. Genetic testing revealed homozygous *Npc1* mutation in both patients (c.2974G>T; p.G992W) (Macias-Vidal, Giros et al., 2014).

### Histology

Eight weeks old animals have been transcardially perfused with 4% PFA/0.1 M PBS followed by 2 h post fixation with 4% PFA/0.1 M PBS. Postnatal day 7 (P7) brains were fixed for 6 h in 4% PFA/0.1 M PBS. Free floating 30 μm sagittal brain sections have been permeabilized and blocked for 1 h in PBS/0.5% Triton X-100/5% normal goat serum (NGS). Next, samples have been incubated O/N at 4°C with primary antibody solution in blocking buffer. After washing 3 times with PBS/0.2% Triton X-100, corresponding goat secondary antibodies (Alexa Fluor, Invitrogen) have been diluted 1:500 in PBS/0.2% Triton X-100/2% NGS and incubated for 1 h at RT. Nuclear staining (Hoechst 1:2000, Invitrogen) or myelin dye (Fluoromyelin red 1:300, Invitrogen) were incubated together with secondary antibodies. Sections have been washed twice with PBS/0.2% Triton X-100 and once with PBS before being mounted using Fluoromount (Sigma Aldrich). Imaging has been performed on at least 3 mice per genotype and time point using a Leica SP5 confocal microscope. Primary antibodies and dilutions were as follows: Calbindin (1:500, Swant); CD68 (1:500, AbDserotec); Iba1 (1:300, Wako); NeuN (1:500, Millipore); CNPase (1:300, Abcam) and Synaptophysin (1:500, Abcam).

### Primary microglia

After the brain had been dissected, cerebellum, olfactory bulb and brain stem were removed. The remaining tissue (cerebrum) was dissociated using the Neural Tissue Dissociation Kit (P) (Miltenyi Biotec) according to the manufacturer’s protocol. To positively select microglial cells, cell suspension was incubated with CD11b microbeads (Miltenyi Biotec) and pulled down using a MACS separation column (Miltenyi Biotec). Purity of microglial fraction was tested by western blot using specific microglial, neuronal, astrocytic and myelin markers as described below (western blot analysis). For mass spectrometry analysis, microglial fraction was washed twice with PBS and subsequently analyzed, without culturing (acutely isolated). For cell culture, isolated microglia were resuspended and plated in DMEM/F12 (Gibco) supplemented with 10% Fetal Bovine Serum (FBS, Sigma Aldrich) and 1% PenStrep (Gibco). For adult microglia, medium was supplemented with 10 ng/ml GM-CSF (R&D System). To analyze cholesterol, microglia from WT and *Npc1*^*-/-*^ animals were plated onto glass coverslips and cultured for 5 days *in vitro* (5DIV). Cells were fixed with 4% PFA/sucrose for 15 min at RT. After permeabilization with PBS/0.1% Triton X-100, cells have been incubated for 1 h in blocking solution (2% BSA/2% FBS/0.2% fish gelatin). Afterwards, primary antibody against CD68 (1:500, AbD Serotec) was incubated for 1 h at RT. After washing, cells were incubated with a goat anti rat Alexa Fluor secondary antibody (1:500, Invitrogen) together with a cholesterol binding dye (Filipin 100 μg/ml, Sigma Aldrich) (Friend & Bearer, 1981) for 1 h at RT. Cells were washed, mounted using Fluoromount and analyzed by confocal microscopy.

### Isolation of peripheral blood monocyte-derived macrophages

Blood samples (20 ml) from clinically affected homozygous *Npc1* mutation carriers and healthy donors were collected. Negative selection of peripheral blood monocyte-derived macrophages was performed by incubating full blood for 1 h at RT with RosetteSep Human Monocyte Enrichment Cocktail (StemCell Technologies). An equal volume of washing buffer (D-PBS/2% FBS/1mM EDTA) was added to each sample and layer of macrophages was separated from red blood cells and plasma by centrifugation on a Ficoll gradient (800 x g for 15 min, GE Healthcare). Potential residues of red cells were eliminated by incubating cell pellets with ACK lysis buffer (Gibco) for 3 min at RT. Lysis buffer was quenched with 40 ml of washing buffer and cells were centrifuged at 300 x g for 7 min. Cell pellets were resuspended and plated in macrophage complete medium (RPMI1640/10% FBS/1% PenStrep/1x Pyruvate/1x NEAA) supplemented with 50 ng/ml hM-CSF (Thermo Scientific). After 48 h, 50 ng/ml of fresh hM-CSF was re-added. At 5DIV, media have been discarded and adherent cells have been washed once in PBS, incubated for 3 min at RT with Versene (Lonza) and scraped in 5 ml macrophage complete medium for further analysis.

### Sample preparation for mass spectrometry

The microglia enriched fractions or human macrophages were lysed in 200 µl of STET lysis buffer (50 mM Tris/150 mM NaCl/2 mM EDTA/1% Triton X-100, pH 7.5) and incubated 15 min on ice with intermediate vortexing. The samples were centrifuged for 5 min at 16,000 x g at 4°C to remove cell debris and undissolved material. The supernatant was transferred to a fresh protein LoBind tube (Eppendorf) and the protein concentration was estimated using the Pierce 660 nm protein assay (ThermoFisher Scientific). A protein amount of 15 µg was subjected to tryptic protein digestion using the filter aided sample preparation protocol (FASP) (Wisniewski, Zougman et al., 2009) using Vivacon spin filters with a 30 kDa cut-off (Sartorius). Briefly, proteins were reduced with 20 mM dithiothreitol and free cysteine residues were alkylated with 50 mM iodoacetamide (Sigma Aldrich). After the urea washing steps, proteins were digested with 0.3 µg LysC (Promega) for 16 h at 37°C followed by a second digestion step with 0.15 µg trypsin (Promega) for 4 h at 37°C. The peptides were eluted into collection tubes and acidified with formic acid (Sigma Aldrich). Afterwards, proteolytic peptides were desalted by stop and go extraction (STAGE) with self-packed C18 tips (Empore) (Rappsilber, Ishihama et al., 2003). After vacuum centrifugation, peptides were dissolved in 2 µl 0.1% formic acid (Biosolve) and indexed retention time peptides were added (iRT Kit, Biognosys).

### LC-MS/MS analysis

For label free protein quantification (LFQ), peptides were analysed on an Easy nLC 1000 or 1200 nanoHPLC (Thermo Scientific) which was coupled online via a Nanospray Flex Ion Source (Thermo Sientific) equipped with a PRSO-V1 column oven (Sonation) to a Q-Exactive HF mass spectrometer (Thermo Scientific). An amount of 1.3 µg of peptides was separated on in-house packed C18 columns (30 cm x 75 µm ID, ReproSil-Pur 120 C18-AQ, 1.9 µm, Dr. Maisch GmbH) using a binary gradient of water (A) and acetonitrile (B) supplemented with 0.1% formic acid (0 min, 2% B; 3:30 min, 5% B; 137:30 min, 25% B; 168:30 min, 35% B; 182:30 min, 60% B) at 50°C column temperature.

Data dependent acquisition (DDA) was used for LFQ. Full MS scans were acquired at a resolution of 120,000 (m/z range: 300-1400; AGC target: 3E+6). The 15 most intense peptide ions per full MS scan were selected for peptide fragmentation (resolution: 15,000; isolation width: 1.6 m/z; AGC target: 1E+5; NCE: 26%). A dynamic exclusion of 120 s was used for peptide fragmentation.

### Mass spectrometry data analysis and LFQ

The data from mouse microglia was analyzed with the software Maxquant, version 1.6.3.3 (maxquant.org, Max-Planck Institute Munich) (Cox, Hein et al., 2014). The MS data was searched against a reviewed canonical fasta database of *Mus musculus* from UniProt (download: November the 5th 2018, 17005 entries). Trypsin was defined as a protease. Two missed cleavages were allowed for the database search. The option first search was used to recalibrate the peptide masses within a window of 20 ppm. For the main search peptide and peptide fragment mass tolerances were set to 4.5 and 20 ppm, respectively. Carbamidomethylation of cysteine was defined as static modification. Acetylation of the protein N-terminal as well as oxidation of methionine were set as variable modifications. The false discovery rate for both peptides and proteins was adjusted to less than 1%. The “match between runs” option was enabled with a matching window of 1.5 min. LFQ of proteins required at least one ratio count of unique peptides. Only unique peptides were used for quantification. Normalization of LFQ intensities was performed separately for the age groups because LC-MS/MS data was acquired in different batches. The protein LFQ reports of Maxquant were further processed in Perseus (Tyanova, Temu et al., 2016). The protein LFQ intensities were log2 transformed and log2 fold changes were calculated between NPC-deficient and WT samples separately for the different age groups, mouse models and patients. Only proteins with a consistent quantification for all three samples per age group were considered for statistical testing. An unpaired Student’s t-test with two-tailed distribution was applied to evaluate the significance of proteins with changed abundance. Additionally, a permutation based false discovery rate (FDR) estimation was used (Tusher, Tibshirani et al., 2001).

The proteomic data was further analyzed through the use of Ingenuity Pathway Analysis (IPA, QIAGEN Inc., https://www.qiagenbioinformatics.com/products/ingenuity-pathway-analysis). Standard settings were used for the analysis. Proteins with a log2 fold-change more than ± 0.5 and a p-value less than 0.05 were defined as protein regulation thresholds.

The data from human macrophages was analyzed with Maxquant version 1.6.3.3 using the same settings searching against a reviewed fasta database of *Homo sapiens* from UniProt including isoforms (download: December the 17th 2018, 42432 entries).

### Western blot analysis

Acutely isolated microglia or human macrophages were lysed in RIPA buffer containing 1% Triton X-100 and supplemented with protease and phosphatase inhibitors (Sigma Aldrich). Lysate protein content was quantified using Bradford assay (Biorad) according to manufacturer’s protocol. At least 10 μg per sample were separated on a bis-tris acrylamide gel followed by western blotting either on a PVDF or nitrocellulose membrane (Millipore) using the following antibodies: Iba1 (1:1000, Wako); GFAP (1:1000, Dako); Tuj1 (1:1000, Covance); CNPase (1:1000, Abcam); NPC1 (1:1000, Abcam); NPC2 (1:1000, Sigma Aldrich); LAMP1 (1:1000, Sigma Aldrich); Cathepsin B (1:2000, R&D System); Cathepsin D (1:1000, Novus Biologicals); mouse CD68 (1:1000, AbDserotec) ; mouse CD63 (1:1000, Abcam); human CD63 (1:500, Santa Cruz); Grn (clone 8H10, (Gotzl et al., 2014)); Trem2 (clone 5F4, (Xiang et al., 2016)); ApoE (1:1000, Millipore); Itgax (1:1000, LSBio); Cath L (1:1000, R&D System); Tmem119 (1:1000, Abcam); P2ry12 (1:1000, Abcam); mTor (1:1000, Cell Signaling); Rab7a (1:1000, Cell Signalling); LC3 (1:1000, Cell Signalling) and Plp1 (1:1000, Abcam). Blots were developed using horseradish peroxidase-conjugated secondary antibodies (Promega) and the ECL chemiluminescence system (GE Healthcare). An antibody against calnexin (1:1000, Stressgen) was used as loading control. For each microglial genotype and time point (P7 and 8 weeks) at least 2 independent cell isolations were used.

### Ex vivo Aβ plaque clearance and myelin phagocytic assay

To functionally characterize myeloid cells (primary microglia and human macrophages), we adapted a phagocytic assay established by Bard et al. (Bard et al., 2000). Briefly, a 10 μm brain section from an AD mouse model (APPPS1) was placed on a poly L-lysine coated glass coverslip. In order to stimulate recruitment of microglia to the plaque site, brain sections were incubated for 1 h at RT with an antibody against human Aβ (5 μg/ml 6E10, BioLegend for microglia and 3 μg/ml 2D8 (Yamasaki, Eimer et al., 2006) for macrophages). Isolated cells were plated at a density of 3×10^5^ (microglia) or 2,5×10^5^ (macrophages) cells/coverslip and incubated either for 5DIV (microglia) or 1DIV (macrophages) in corresponding culturing medium. Next, samples were fixed with a 4% PFA/sucrose solution for 15min at RT. Immunostainings were performed as described above using an antibody against myeloid cells (anti-CD68 for microglia and anti-CD45 for macrophages) and Aβ plaque (3552, (Yamasaki et al., 2006)). Fibrillar Aβ (plaque core) was visualized using Thiazine red (ThR 2 μM, Sigma Aldrich) added into the secondary antibody solution. To evaluate the phagocytic capacity of myeloid cells, plaque coverage (ThR signal area for microglia or 3552 for macrophages) was quantified, comparing brain section incubated with cells with a consecutive brain section where no cells were added. Quantification was done using a macro tool in ImageJ (NIH), applying to 10x 16-bit tail scan picture of the whole coverslip with a threshold algorithm (OTSU) and measuring particles with a pixel size from 5 to infinity. Each experimental group was tested in at least 3 independent experiments.

Similarly, we applied this assay to evaluate the capability of P7 microglia to uptake and digest myelin from the APPPS1 brain section. In order to evaluate myelin level after incubation with microglia, we used Fluoromyelin dye (1:300, Invitrogen) added together with the secondary antibody solution. CD68 was used to visualize microglia. Each experimental group was tested in at least in 3 independent experiments.

### In vitro myelin phagocytic assay

Purified myelin was isolated as previously described (Cantuti-Castelvetri et al., 2018). Briefly, 8 weeks old C57BL/6J mouse brains were homogenized by sonication in 10 mM HEPES/5 mM EDTA/0.3 M sucrose/protease inhibitors. The homogenate was layered on a gradient of 0.32 M and 0.85 M sucrose in 10 mM HEPES/5 mM EDTA (pH 7.4) and centrifuged at 75,000 x g for 30 min with a SW41 Ti rotor (Beckman Coulter). The myelin fraction was isolated from the interface, and subjected to 3 rounds of osmotic shock in sterile ultrapure water and centrifuged at 75,000 x g for 15 min. The resulting pellet was subjected to the same procedure to obtain a pure myelin fraction. The yield of myelin was calculated by measuring the total amount of protein with the Bradford assay (Biorad). For the fluorescence labeling of myelin, a PKH67 Green Fluorescent Cell Linker Mini Kit was used (Sigma Aldrich).

Primary microglia from P7 pups and human macrophages have been isolated and cultured as described above. At 5DIV, cells were fed with purified green fluorescently labeled myelin at a concentration of 50 μg/ml. At 6 h, myelin was washed out and cells were fixed at 24/48/72 h with a 4% PFA/sucrose solution for 15 min. Immunostaining of fixed cells was performed as described above using antibodies against CD68 and Perilipin 2 (1:200, Progen) for microglia and CD45 for patient macrophages. Microglial lipid droplets were visualized using Nile red staining kit (Abcam) according to manufacturer’s protocol. In order to monitor myelin trafficking during phagocytosis, primary P7 microglia were incubated with cholera toxin (CTX) subunit B Alexa Fluor(tm) 647 Conjugate (0,5 μg/ml, Thermo Fisher Scientific) and Red DQ-bovine serum albumin (DQ-BSA; 10 μg/ml, Invitrogen) for 30 min at 37°C in microglial culturing medium. Cells were pulsed for 15 min with green fluorescently labeled myelin (10 μg/ml) and fixed after 1 h. Three independent cell preparations were treated and analyzed by confocal microscopy.

### EM analysis

The microglial pellet was preserved throughout all fixation, contrasting and infiltration steps. Cells were fixed for 30 min in 2% PFA/2.5% glutaraldehyde (EM-grade, Science Services) in 0.1 M sodium cacodylate buffer (pH 7.4) (Sigma Aldrich), washed three times in 0.1 M sodium cacodylate buffer before postfixation in reduced osmium (1% osmium tetroxide (Science Services)/0.8% potassium ferrocyanide (Sigma Aldrich) in 0.1 M sodium cacodylate buffer). After contrasting in aqueous 0.5% uranylacetate (Science Services), the pellet was dehydrated in an ascending ethanol series, infiltrated in epon (Serva) and cured for 48 h at 60°C. 50 nm ultrathin sections were deposited onto formvar-coated copper grids (Plano) and postcontrasted using 1% uranyl acetate in water and ultrostain (Leica). Transmission Electron Microscopy micrographs were acquired on a JEM 1400plus (JEOL) using the TEMCenter and tile scans with the ShotMeister software packages (JEOL), respectively.

### Statistical analysis

For comparison between two groups, unpaired Student’s t-test with two-tailed distribution was used. Data are represented as mean ± standard error of the mean (SEM). A value of p < 0.05 was considered significant (*p < 0.05; **p < 0.01; ***p < 0.001).

## Acknowledgements

We thank Baccara Hizli for helping with human blood sample collection and Anna Berghofer for excellent technical assistance. APPPS1 mice were kindly provided by Mathias Jucker (Hertie-Institute for Clinical Brain Research, University of Tübingen). The authors thank Anja Capell, Dieter Edbauer, Christian Haass, Matthias Prestel and Michael Willem for critically reading the manuscript. This work was supported by the NCL Foundation and the Deutsche Forschungsgemeinschaft (DFG, German Research Foundation) within the framework of the Munich Cluster for Systems Neurology (EXC 2145 SyNergy). S.A.S was supported by a LMU Excellence Program for Clinician Scientists, Verum-Stiftung and the Ara Parseghian Medical Research Foundation.

## Author Contributions

S.T. and A.C. designed and supervised the study and wrote the manuscript with the input from all co-authors. A.C. performed human and animal experiments including histology, microglia and human macrophage experiments, target validation and functional studies. S.H. contributed to conceptual design. L.D. and L.V. contributed to histological data. L.D. and L.S.M. assisted in isolation of primary microglia. M.Sch. performed EM analysis of isolated microglia. L.C.C and M.S. contributed to the design of the *in vitro* myelin phagocytic assay. S.A.M., J.K. and S.F.L. performed the proteomic analysis. T.B.E., S.A.S. and M.Str. recruited NPC patients and control individuals and delivered study samples.

## Expanded View Figure Legends

**Figure EV1. Pronounced microgliosis in symptomatic *Npc1***^***-/-***^ **mice. A-C** Immunostaining of cerebellum (**A**), cortex (**B**) and hippocampus (**C**) of WT and *Npc1*^*-/-*^ brain sections with antibodies against neuronal markers (green) calbindin (**A**, Purkinje cells), NeuN (**B-C**) and lysosomal microglial marker CD68 (red). Low magnification (10x, upper panels) analysis of *Npc1*^*-/-*^ brain shows evident Purkinje cell loss in the cerebellum while no significant neuronal loss is observed within the cortex and hippocampus. High magnification images (100x, lower panels) show amoeboid microglial morphology in *Npc1*^*-/-*^ brains. Hoechst was used for nuclear staining (blue). Scale bars: 250 μm (10x, upper panels) and 25 μm (100x, lower panels).

**Figure EV2. Quality control for microglial isolation from 8 weeks old mouse brain tissue using MACS separation system.** Total protein lysates from microglia enriched and microglia depleted fractions were analyzed via western blot analysis for cell specific markers of microglia (Iba1), neurons (Tuj1), astrocytes (GFAP) and oligodendrocytes (CNPase). Calnexin was used as a loading control.

**Figure EV3. Canonical pathway analysis using the IPA web-tool.** The ten most significantly affected pathways for 8 weeks and P7 *Npc1*^*-/-*^ microglia are represented as bar charts plotting the negative transformed log10 p-value.

**Figure EV4. Western blot validation of microglial MS data from 8 weeks old mice.** Representative immunoblots of acutely isolated WT and *Npc1*^*-/-*^ microglia show increased levels of CD63 and NPC2 and alteration in proteins involved in autophagy (mTOR, p70S6K, RAB7a and LC3I and II). For each protein investigated, microglia coming from 2 independent mice per genotype have been analyzed. Calnexin level was used as loading control. * indicates unspecific bands.

**Figure EV5. Microgliosis precedes neuronal loss in P7 *Npc1***^***-/-***^ **mice.** Immunostaining of hippocampus from P7 WT and *Npc1*^*-/-*^ mouse brains with antibodies against neuronal marker NeuN (green) and lysosomal microglial marker CD68 (red). Hoechst was used for nuclear staining (blue). At P7 we could not detect overt neurodegeneration in the hippocampal region of *Npc1*^*-/-*^ mice, as judged by preserved neuronal architecture, but we could already detect increased CD68 immunoreactivity (10x, upper panels). Imaging analysis confirmed that amoeboid *Npc1*^*-/-*^ microglial morphology (100x, lower panels) is an early pathological hallmark. Scale bars: 250μm (10x, upper panels) and 25μm (100x, lower panels).

**Figure EV6. Microglia show an increased myelin uptake in both pre-symptomatic and symptomatic *Npc1***^***-/-***^ **mice. A** Immunostaining of brain sections from hippocampus of 8 weeks old *Npc1*^*-/-*^ mice shows myelin accumulation (Fluoromyelin, green) within late endosomal/lysosomal compartment (CD68, red). Hoechst is used for nuclear staining (blue). Scale bar: 50 μm. **B** Increase of myelin protein PLP1 could be detected in acutely isolated microglia from symptomatic (8 weeks) and pre-symptomatic (P7) *Npc1*^*-/-*^ mice compared to WT.

**Figure EV7. P7 *Npc1***^***-/-***^ **microglia efficiently uptake myelin but fail in its turnover.** Representative images of an *ex vivo* myelin phagocytic assay where acutely isolated WT and *Npc1*^*-/-*^ microglia were plated onto an APPPS1 brain cryosection and assessed for myelin uptake. Microglial lysosomes were stained with an antibody against CD68 (green) while myelin was visualized using Fluoromyelin (red). Nuclei were labeled using Hoechst (blue). In contrast to intact white matter tracts visualized by Fluoromyelin upon addition of WT microglia, white matter tracts upon addition of *Npc1*^*-/-*^ microglia were disrupted, indicating increased myelin uptake. Uptaken myelin accumulated within CD68 positive compartment in *Npc1*^*-/-*^ microglia, suggesting possible impairments in myelin turnover. Scale bar: 50 μm.

**Figure EV8. Microglial impairment in myelin turnover and lack of lipid droplet staining in *Npc1***^***-/-***^ **microglia.** Primary microglia isolated from P7 WT and *Npc1*^*-/-*^ mice were fed with fluorescently labeled myelin (green) and analyzed after 72 h. In order to confirm the identity of lipid vesicles forming as a result of myelin turnover, we stained microglia with Nile red (red) to visualize lipid droplets. Boxed regions are enlarged in right panels and reveal co-localization between fluorescently labeled myelin vesicles and Nile red, supporting myelin turnover and lipid droplet formation in WT microglia. In *Npc1*^*-/-*^ microglia, Nile red mainly stained myelin deposits accumulating in late endosomes/lysosomes, instead of lipid droplets at the cell periphery, confirming impairment in myelin turnover. Scale bars: 10 μm.

## Expanded View Table Legends

**Table EV1. MS analysis of microglia isolated from symptomatic *Npc1***^***-/-***^ **mice.** Table displays all significantly changed proteins in the *Npc1*^*-/-*^ microglial proteome versus WT, average of protein LFQ intensity ratio, log2 conversion are corresponding p-values. Additionally, a permutation based false discovery rate (FDR) estimation was applied and significantly changed proteins were highlighted (+).

**Table EV2. MS analysis of microglia isolated from pre-symptomatic *Npc1***^***-/-***^ **mice.** Table displays all significantly changed proteins in the *Npc1*^*-/-*^ microglial proteome versus WT, average of protein LFQ intensity ratio, log2 conversion and corresponding p-values. Additionally, a permutation based false discovery rate (FDR) estimation was applied and significantly changed proteins were highlighted (+).

**Table EV3. MS analysis of human macrophages from NPC patients.** Table displays all identified proteins in the proteome of peripheral blood-derived macrophages from NPC patients versus a healthy control. Individual protein LFQ ratios and corresponding log2 conversion are shown separately for patient one (17 years old) and patient two (24 years old).

