## Supplementary figures and images for "Loss of NPC1 enhances phagocytic uptake and impairs lipid trafficking in microglia"

### Supplemental files

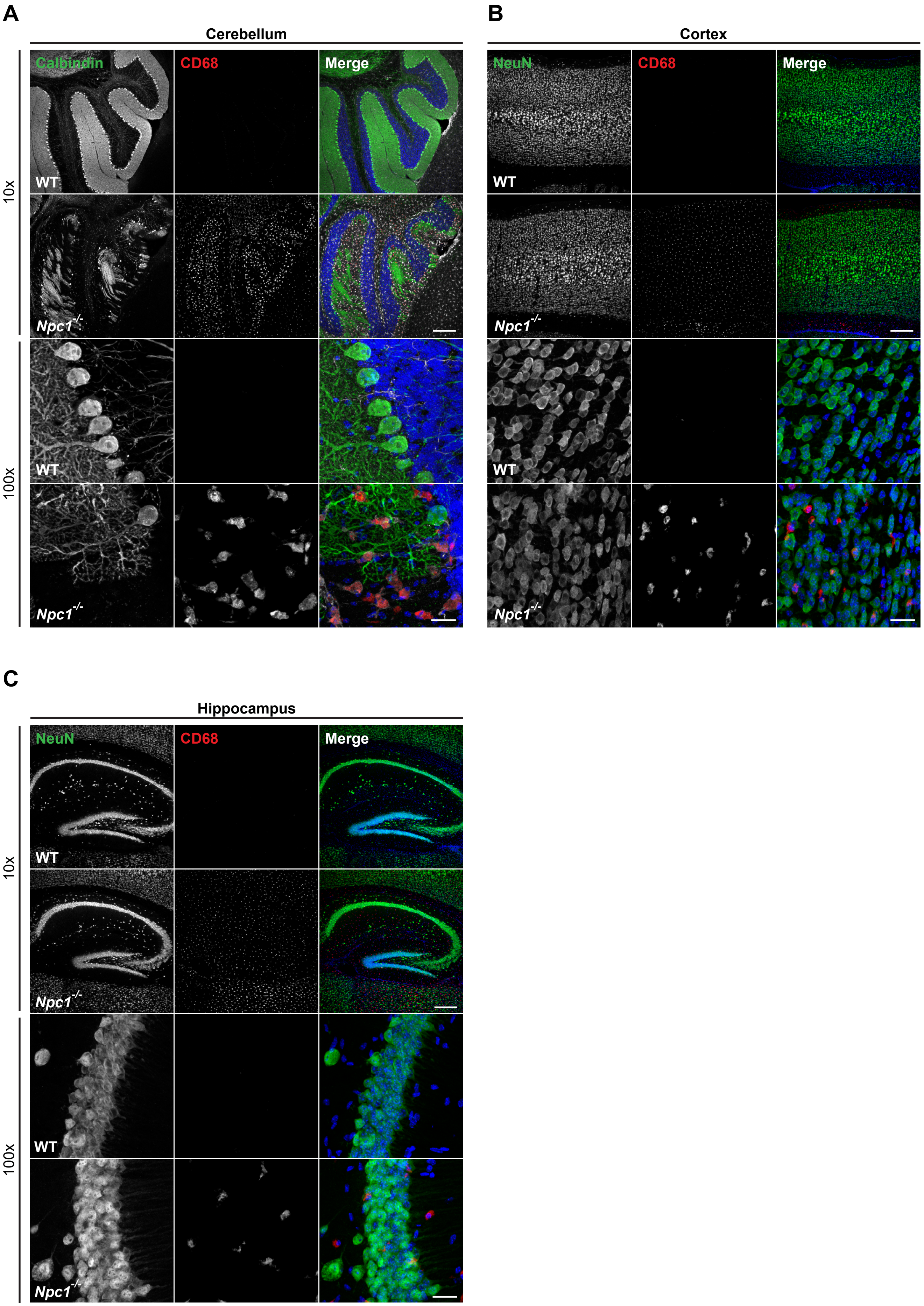

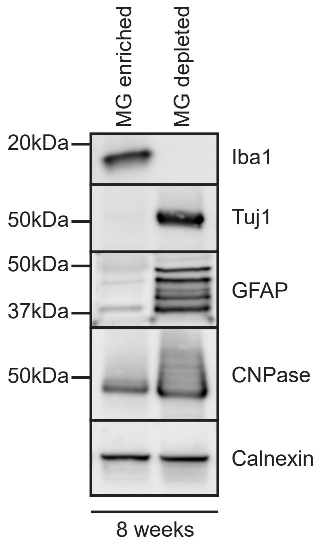

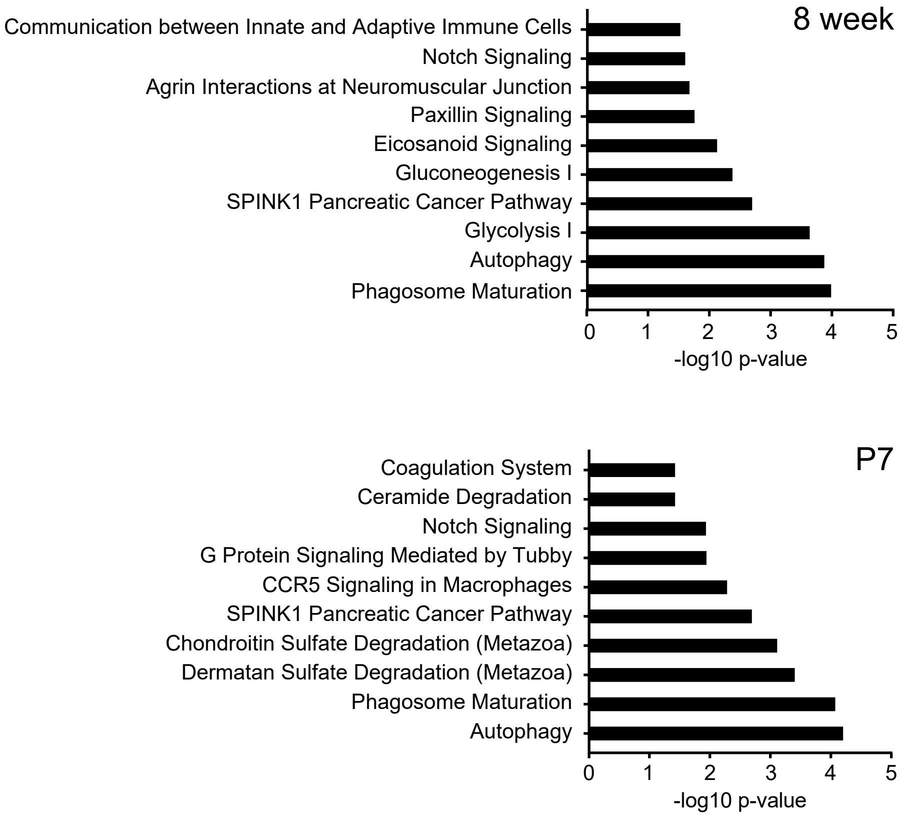

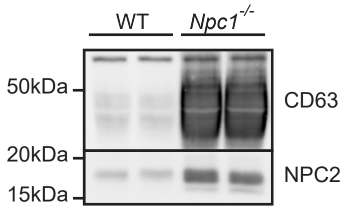

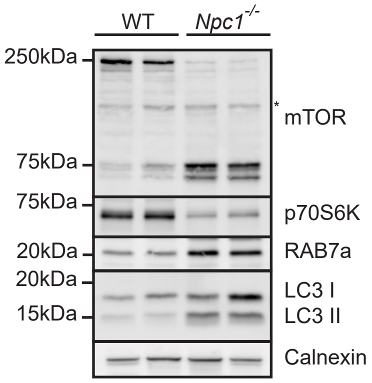

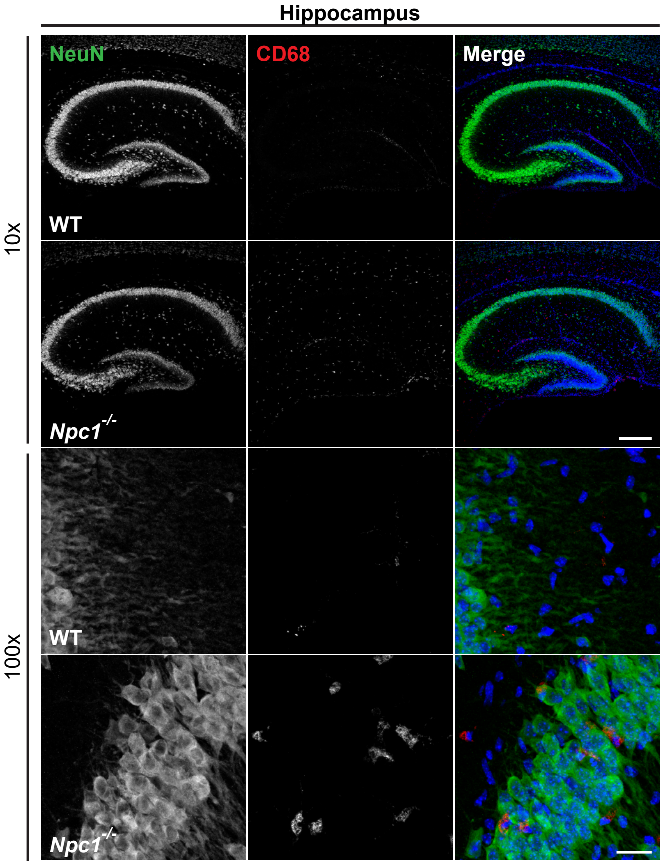


Fig EV5

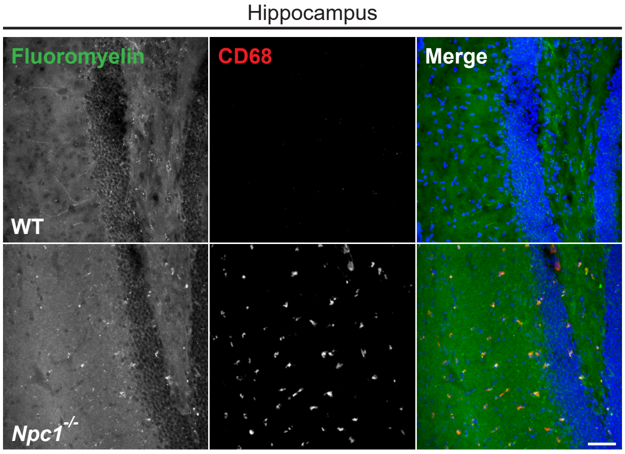

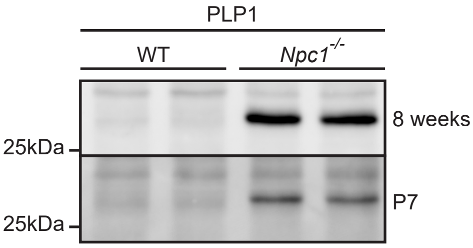

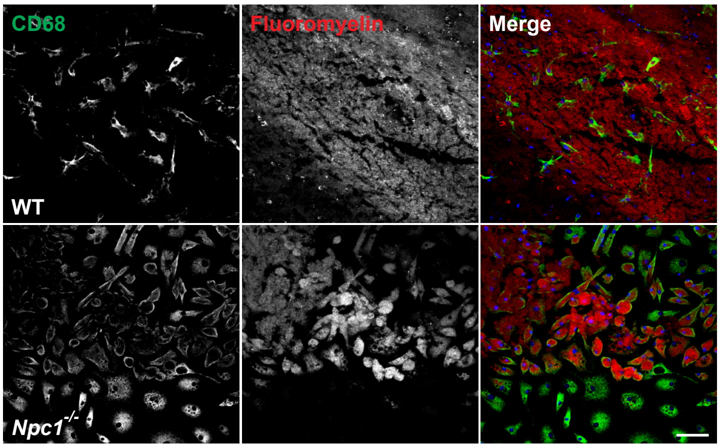


Fig EV7

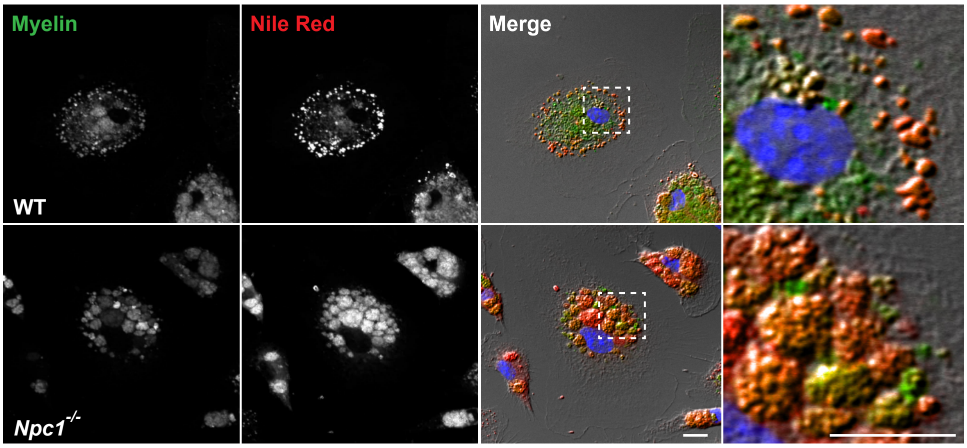


Fig EV8
